# Separable neural population representations are constructed from mixed single neuron selectivity in the mouse early visual system

**DOI:** 10.1101/2025.08.25.672226

**Authors:** Juan Santiago Moreno, Nicholas Garcia, Daniel J. Denman

## Abstract

Both sensory and non-sensory brain regions receive mixed inputs from single neurons which require decomposition and integration before proceeding through a processing hierarchy. Whether mixed input signals are used in biological neural networks to derive pure single neuron representations, or distributed as new population representations from mixed single neurons, is not clear. In this study, we measured the distribution of single neuron hue and luminance tuning in the dorsolateral geniculate nucleus (dLGN) and primary visual cortex (V1) of mice, as well as the information about and structure of hue and luminance representations in populations of hundred of simultaneously sampled neurons. We compare single neuron and population encoding to null models expected for random integration and extraction of pure categorical single neuron representation. Using both univariate and multivariate regression techniques, we consistently noted that tuning for hue and luminance, rather than clustering into categorical response structures, formed uniform distributions. While the distribution of single neuron selectivity varied across the thalamocortical circuit, we found no evidence of categorical tuning organization emerging in the hierarchy. Nevertheless, populations contained complete information, in either high-dimensional linear representations or low-dimensional non-linear representations. In summary, we find that as early as primary sensory cortex and thalamus single neurons that have mixed selectivity for hue and luminance form a high dimensional representation of those variables, which can be non-linearly embedded in multiple separable representations.

## INTRODUCTION

Many brain areas are thought to compute by combining multiple types of input information to create single neuron specificity. Examples can be found in both sensory [1–3] and non-sensory systems [4, 5]. At the same time, a growing body of evidence points to “mixed” selectivity within single neurons across the brain - within [6–9] and across [10–12], modalities. If populations of single neurons with mixed signals represent categories of information, and how such population representations are maintained or transformed across areas, are unresolved questions in neuroscience. Further, insights into how neural systems are able to use a limited set of resources to form general representations that enable flexible behavior will inform computation from sensory to cognitive systems [13].

In the early visual system, color, specifically hue, is thought to be encoded through parallel channels of color-opponent single neurons originating in the primate retina [14, 15], and these parallel representations persist in downstream areas through V1 [16–19]. Although hue and luminance are inherently mixed as qualities of the same photons, they can be made perceptually distinguishable by distinct signaling mechanisms in the retina. In primates, parallel color processing pathways are maintained by single neurons at least through the primary visual cortex (V1) [16, 20, 17, 21], though other studies have demonstrated less rigid categories of single neuron selectivity, roughly divided into luminance-preferring, color-luminance, or color-preferring, with the color-luminance population having a broad distribution of color preference [18, 22]. In the mouse, many cones co-express the two available opsins and a subpopulation express only the short opsin [23]. As such, variations in wavelength *or* luminance can drive changes in individual cones containing both types of cone opsin expression [14, 24]. Distinct luminance and hue signaling populations must be generated and maintained over a successive recombination that give rise to form representation centrally [25] and perceptually [26, 27]. Given the mixed nature of these signals from their origins, it is unknown if or how these hue and luminance signals are disambiguated by the downstream sampling strategies in central visual neural populations or how such transformations may occur within and between visual areas.

Here, we measure the relationship between mixed single neuron selectivity and the resultant population representations of hue and luminance in the mouse early visual system. We survey single neuron selectivity at a large scale and quantify the representations in the simultaneous activity of multiple populations, comparing the observed hue and luminance single neuron selectivity to quantitative null models of categorical and random selectivity, finding few categorical neurons in the population of “mixed” single neurons. In addition to exploring the distribution of chromatic signals in single neurons, we measure decodable information about each signal in these mixed populations, and the low-dimensional structure of the population representation.

## RESULTS

We sought to measure the population representation of opsin-specific luminance, overall luminance, and hue in the mouse dorsolateral geniculate nucleus (dLGN) and primary visual cortex (V1). In order to generate statistical expectations of categorical and randomly distributed spectral tuning representations, we first created two spiking models to simulate responses to arbitrary stimuli. In those models, single neuron tuning was either *purely selective* or *randomly selective* with respect to opsin content, respectively (Figure 1A). Because luminance is produced by additive opsin signals, the axis of categorical luminance selectivity indicates equal increments or decrements in M and S-opsin contrast. Likewise, because hue is produced by contrasting opsin signals, the axis of categorical hue selectivity is defined perpendicular to the axis of luminance selectivity (Figure 1B). Any neuron whose tuning deviated from these axes would have mixed selectivity for color and luminance in some relative proportion. To formalize this conceptual model, and generate a set of predictions for observations in the early visual system, we simulated spiking populations (see Materials and Methods) by explicitly defining the distribution of tuning preferences (Figure 1C, top) for each model, categorical (blue box) and random (orange box). We then simulated the response of populations of neurons defined by these models to changes in M and S luminance. The resulting pure single neuron selectivity in the categorical model was distributed along the axes of color and luminance (Figure 1C, bottom). The random model yielded mixed single neuron preferences distributed throughout mouse color space, with a bias towards the origin, indicating an overall reduction in both color and luminance information in single neuron selectivity. While these frameworks yielded strikingly different distributions of single neuron tuning metrics, we note that both produce population-level representations that contain decodable information about each opsin contrast, overall luminance, and hue (Figure 1D). These models therefore provide statistical expectations at potential extremes of representational frameworks for hue and luminance: null models which we can use to compare to observed representation in experimental data in dLGN and V1.

**Figure 1:**
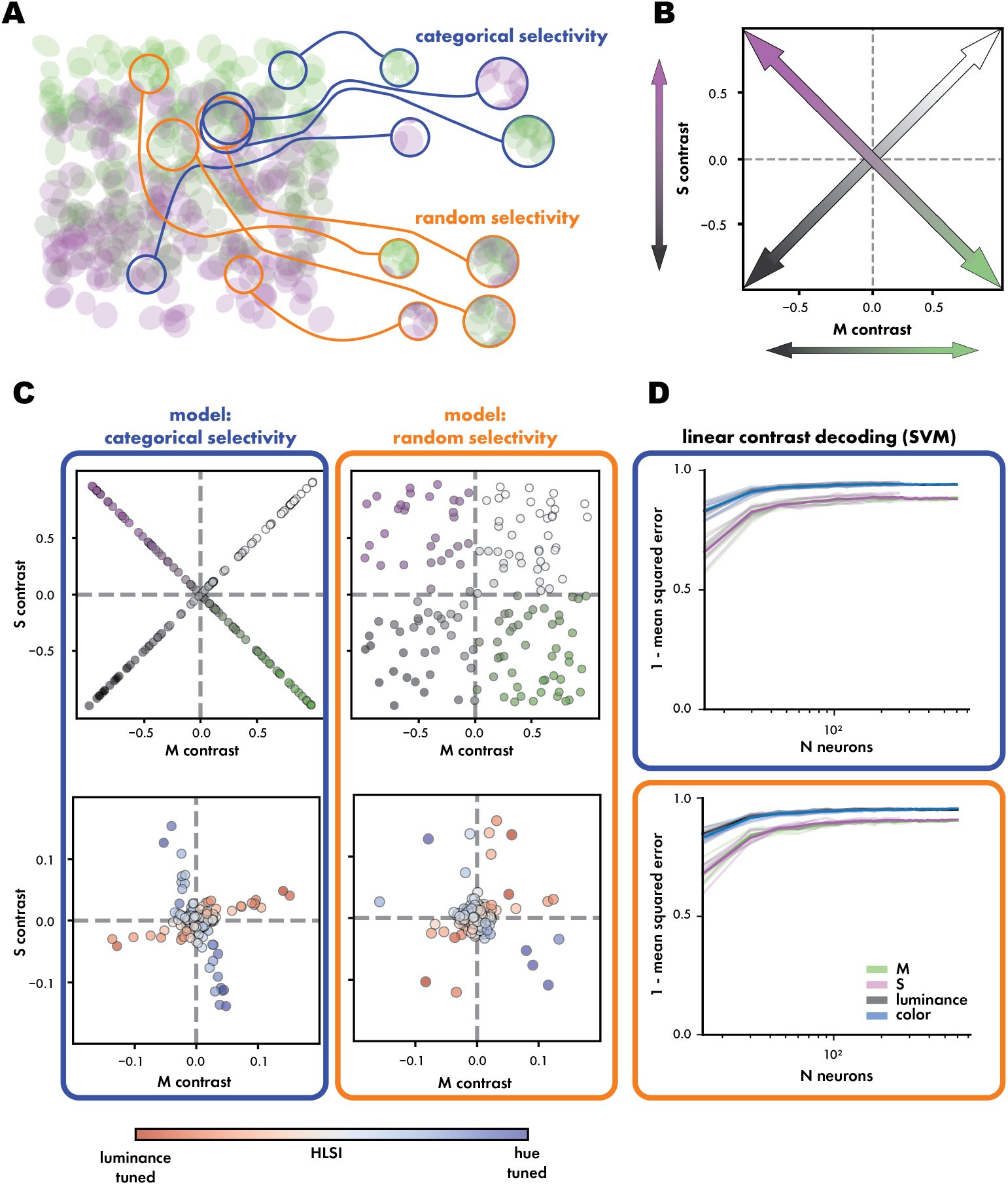
Both categorical and random sampling of the opsin mosaic can yield down-stream single neuron selectivity and population hue and luminance information. (**A**) Conceptual model of downstream convergence of opsin signals via categorical and random selectivity mechanisms. (**B**) Mouse color space (M opsin contrast versus S opsin contrast) with axes of hue (pink and magenta) and luminance (black and white). (**C**) Statistical null models of conceptual models of downstream converge, and the resulting predicted distributions of single neuron selectivity (**D**) Performance of modeled populations linearly decoding stimulus contrasts using support vector regression (SVR). Performance is represented as 1 −mean squared error (MSE) relative to population size.

### Populations of dLGN and V1 neurons are modulated by full-field changes in luminance and hue contrast

To explore how thalamocortical circuits compare to these hypothesized representations for integration of luminance and hue signals *in vivo*, we simultaneously recorded neurons in dLGN and all layers of V1 in awake, head fixed mice (Figure 2A) while presenting a set of chromatic, achromatic, spatially structured, and spatially uniform stimuli (Figure 2B) – luminance flashes (achromatic, uniform), color exchange (chromatic, uniform), drifting gratings (achromatic, structured), and chromatic gratings (chromatic, structured). Luminance flashes and uniform color exchange in stimuli allowed us to measure distinct and complementary components of single neuron tuning in both dLGN and V1. Stimuli were presented over the full visual field using a custom spherical projection apparatus (Figure 2A). Projector LEDs were selected to align to the peak spectral sensitivities of mouse cone opsins (Figure 2C). Using high-density electrophysiology (Neuropixels probes [28, 29]) allowed us to capture the simultaneous activity of 69-815 neurons per recording for a total of 3700 neurons across 8 recordings. Each unit was assigned to a structure or layer within that structure using depth estimation aligned to electrophysiological landmarks (Figure 2D, Figure 2 with Figure S1). Each cortical unit was classified as fast spiking (FS) or regular spiking (RS) based on their waveform morphology and duration (Figure 2E).

**Figure 2:**
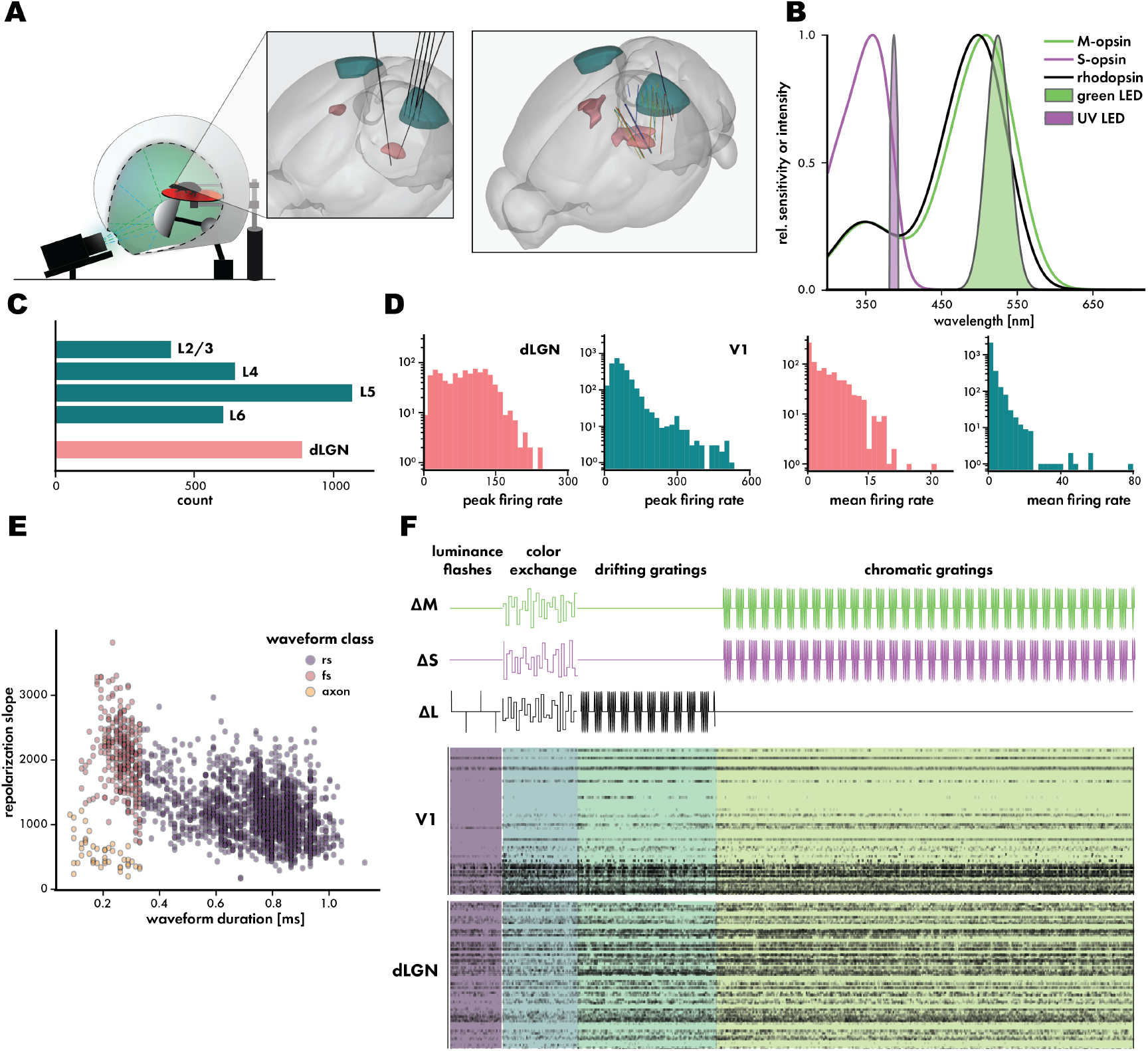
Populations of dLGN and V1 neurons are modulated by luminance and color contrast. (**A**) Illustration of immersive stimulus environment and example probe insertion trajectories alongside a rendering of all probe trajectories from recordings included in this project. (**B**) Mouse cone opsin spectra [65] overlaid with spectra of green and UV projector LEDs. (**C**) Number of neurons recorded from dLGN (light coral) and each V1 layer (dark cyan) across recordings (**D**) Distribution of peak (**left**) and mean (**right**) firing rates in dLGN and V1 across recordings. Further details and alignment to the Allen Institute Common Coordinate Framework [62] to generate these labels is in Figure 2 with Figure S1. (**E**) Scatter distribution of waveform duration versus repolarization slope, which were used to label cortical units as fast-spiking neurons (FS), regular spiking neurons (RS), or axons (**F**) Illustration of the visual stimulus sequence used in our recordings. (**above**) Directly modulated changes in M opsin, S opsin, or luminance contrasts. (**below**) Spiking activity across probe channels inserted into dLGN and V1 during an example recording

### Selectivity for color and luminance contrasts are uniformly distributed in opsin contrast space

Using these population recordings, we considered single neuron selectivity to presented changes in spectral contrast. Previous work has measured chromatic selectivity in mouse dLGN [30, 31] and V1 [32–34, 25, 35], but cortical observations have been limited to superficial layers with inferred spiking, leaving the dynamics, deeper layers, and relationship with dLGN single neuron chromatic selectivity less well characterized. To address this, we used brief, spatially-uniform, anisoluminant changes in color contrast to drive cones across the majority of retinotopy. Rather than measuring neurons’ preferences for each LED, we calibrated stimuli based on their relative stimulation of M and S opsins [36] to generate radially symmetrical M-S opsin contrasts (Figure 3 with Figure S3).

**Figure 3:**
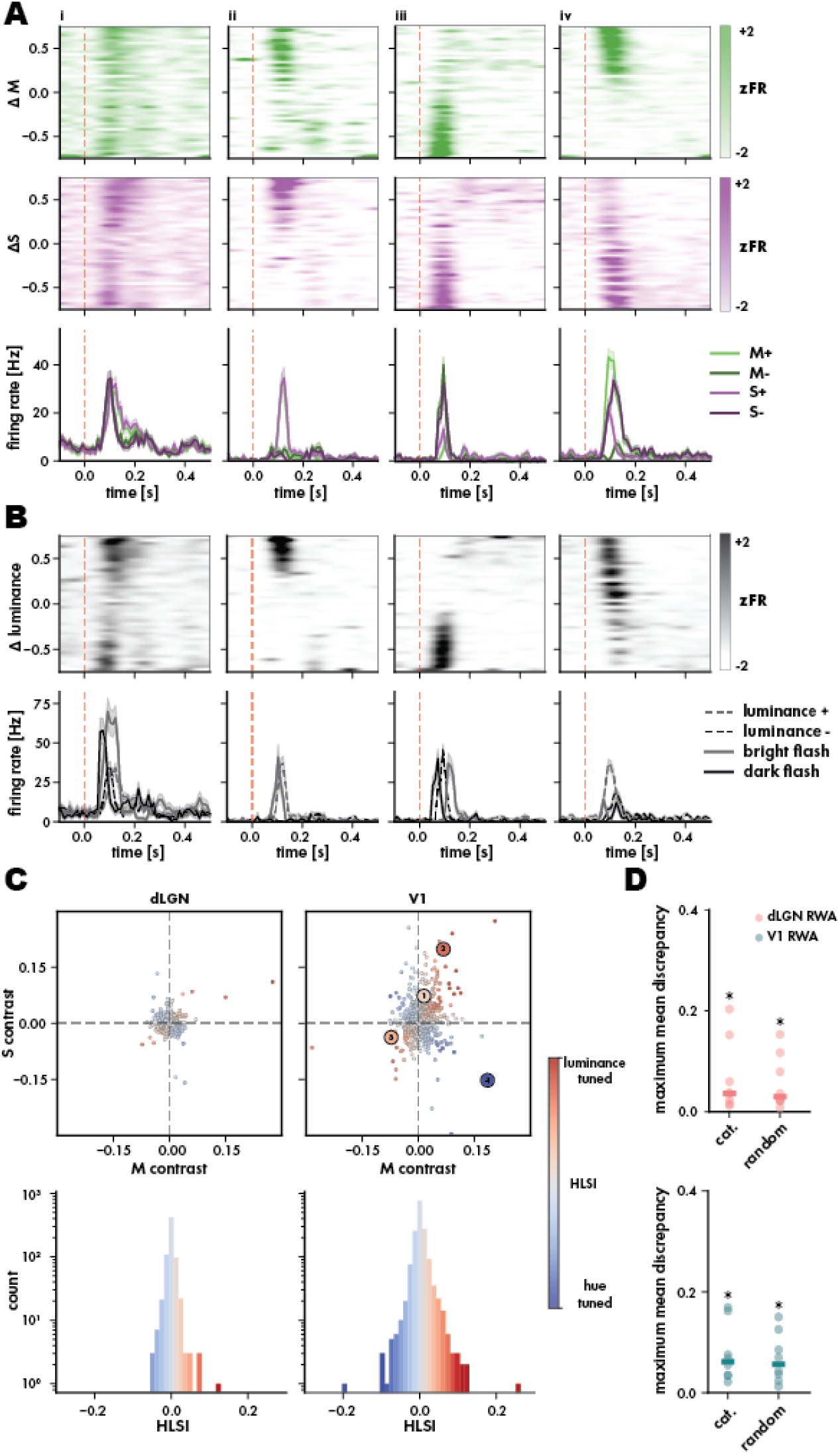
Single neurons display high luminance and color selectivity, but the overall population representation does not support categorical selectivity. (**A**) Example V1 neurons with categorical tuning profiles. Trial responses organized by change in M contrast (**top**) and S contrast (**middle**). Mean trial responses grouped by increment or decrement in opsin contrasts (bottom). From left to right: **i**) unsigned preference for luminance contrast **ii**) luminance-ON **iii**) luminance-OFF **iv**) hue opponent (M-ON/S-OFF) (**B**) Example neurons as in A. Trial responses organized by changes in luminance contrast (**top**). Trial responses condensed by increment or decrement in luminance contrasts compared to same neurons’ responses to full-field luminance flash. (**C**) Scatter plots of RWA for M versus S contrast in LGN and V1, colored by each neuron’s HLSI (**top**). Histograms of HLSI distribution in both regions (**bottom**) (* p¡ 0.05; Wilcoxon rank-sums).

We measured the hue and opsin-specific luminance contrast responses for 1599 V1 neurons and 685 dLGN neurons (n = 8 recordings, n = 8 mice, mean firing rate during stimulus ¿ 0.5Hz). Across both populations, we found diverse response profiles to contrast changes that included several neurons with categorical response profiles (Figure 3A), as might be expected in our categorical model and from the literature [20]. Single neuron response profiles included strong responses to both increments and decrements in M and S contrast, denoting unsigned preference for luminance (Figure 3Ai). Individual neurons also fell along the categorical luminance axis, generating strong responses to increments in luminance (ON cell, Figure 3Aiii) and decrements in luminance (OFF cell, Figure 3Aiii) in the range of each cone opsin. Finally, consistent with previous reports [26, 32, 35] we observed single neurons through our sample with strong hue selectivity (Figure 3Aiv, M-ON/S-OFF). The qualitative category of these responses relative to the total luminance contrast (sum of changes in M and S opsin contrasts) were not meaningfully different from responses driven by full field luminance flashes (compare Figure 3Ai-iv to Figure 3Bi-iv). Notably the M-ON/S-OFF neuron was an overall luminance ON cell due to relatively stronger M-ON responses relative to S-OFF.

Using these responses, we calculated a response weighted average (RWA) for each neuron to changes in each opsin contrast: using the sum of the baseline-corrected response on each trial (labeled by the respective change in M or S contrast) we calculated a weighted average of the contrasts presented [37]. This approach is beneficial for two reasons: 1) compared to some prior studies evaluating chromatic selectivity based on hue’s relative green-UV content, it better reflects the mixed-opsin inputs from M-cones at the retina and as a result, 2) it allows us to simultaneously measure preferences for hue contrast and luminance contrast. Each neuron’s RWA is a point in opsin contrast space (Figure 3C), which facilitates the direct comparison to pure selective and random models (Figure 1). We labeled each neuron with a *hue-luminance selectivity index (HSLI)* which indicated how closely each point aligned to the categorical axes of either hue (negative) or luminance (positive) multiplied by that point’s strength of tuning (distance from the origin). While the examples of categorical tuning (Figure 3A,B) align to their expected axes, the vast majority of neurons are classified in-between hue and luminance tuning (Figure 3C). To quantify distributional uniformity, we computed the normalized entropy of the 2D tuning distribution for dLGN (H = 0.823; range 0–1, where 1 is perfectly uniform) and V1 (H = 0.768). We compared this to a null distribution generated by bootstrap sampling from a 2D uniform distribution. The observed entropies for both distributions were not significantly different from the null expectation under a uniform distribution (p = 1.0 and 1.0 respectively), indicating the distributions were statistically indistinguishable from a uniform set of preferences, with no clustering. Despite notable qualitative differences in the X-shaped vs more uniform shapes of the tuning distributions in the categorical vs random models, we sought to quantify the degree of dissimilarity between the experimental samples and the model distributions. For this, we computed the *maximum mean discrepancy* (MMD, Figure 3D). An MMD of 0 indicates identical distributions and higher MMD values indicate a greater difference in the joint structure (e.g., orientation or multimodality) of the 2D distributions. To avoid larger discrepancies resulting from differences in the overall ranges of the distributions, we scaled distributions before comparison to have a mean of zero and unit variance. We found that the median MMD value across experiments for dLGN and V1 populations were comparable against both models, suggesting similar dissimilarity of both areas from uniform and from random. The MMDs were significantly greater than the size-matched null MMD values (p ¡ 0.05, Wilcoxon rank-sum, Figure 3D). In aggregate these measures indicate both dLGN and V1 have non-random, but also non-categorical distributions (Figure 3D) of luminance and hue selectivity, uniformly covering color space.

While both areas had a statistically uniform distributions of single neurons across color space, we inspected the distributions for potential differences in the representations between V1 and dLGN. When considering opsin contrast tuning independently, V1 had a wider range of tuning than dLGN for both M (dLGN: [−0.073,0.317], V1: [−0.284,0.205])and S opsin (dLGN: [−0.159,0.133], V1: [−0.296,0.274]). dLGN was heavily right-skewed toward negative S contrast, but tuning for M contrast was essentially normally distributed (M: skew = −0.220, S: skew = 7.138). The pattern in V1 was similar, but the S contrast skew was not as strong (M: skew = 0.268, S: skew = 1.113). Using HLSI to compare the the dLGN and V1 distributions as 1-dimensional distributions of hue versus luminance tuning, V1 had a greater spread than dLGN ([−0.050,0.120] and [−0.195,0.256], respectively) and both significantly deviated from normality (Shapiro-Wilk; dLGN = p ¡ 0.001; V1= p ¡ 0.001). Both distributions were strongly right-skewed, indicating a bias toward color tuning, with LGN showing greater skew (skew = 1.87) than V1 (skew = 1.19). Both areas showed very strong kurtosis (dLGN: kurtosis = 16.842, V1: kurtosis = 23.996), indicating a large number of outliers outside of the sharp central peak. These results demonstrate that despite examples of categorical single neuron tuning, both dLGN and V1 have non-random, non-categorical representations; these single neuron distributions are uniformly distributed in both dLGN and V1, but with a marked expansion in the range of spectral contrast selectivity in V1.

### Model-based estimation of single neuron contribution to mixed hue and luminance representations

The RWA metric assesses single neuron response profiles against *independent* individual variables, though we know that hue and luminance depend on each other in visual inputs and opsin contrasts and drive mixed effects through retinal circuits. To find the association between single neurons in populations and multiple variables simultaneously, we used a linear encoding model to estimate the effect size of each contrast type on responses. Reduced-rank regression (RRR) predicts the activity of single neurons in response to changes in contrast that occur during each trial. Using a small set of shared temporal bases learned from the whole population activity, RRR reduces the number of parameters to mitigate overfitting issues. The prediction of time-varying responses specific to each variable of interest (hue and luminance contrasts) provides a more accurate metric of selectivity than trial-averaged metrics which do not distinguish which component of the response is related to each contrast. The resulting selectivity profile is more readily comparable across different stimulus features, has better predictive power, and mitigates issues from unbalanced or missing stimulus conditions compared to traditional trial-averaged approaches [38]. For each trial of the color exchange, we considered a time window of −0.1 to 0.4 seconds relative to stimulus onset, based on the 2Hz frequency of stimulus presentation (Figure 4A).

**Figure 4:**
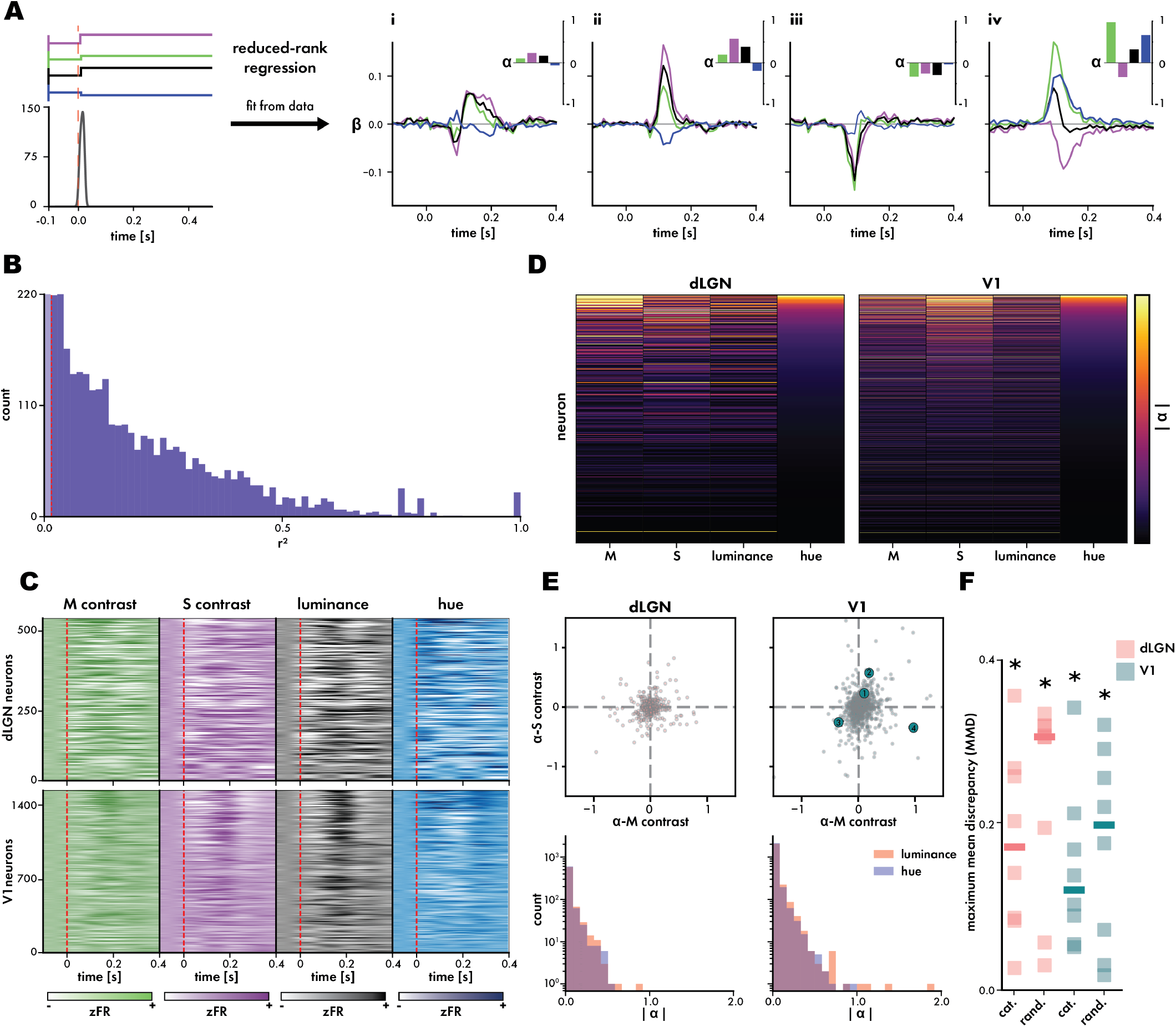
Responses dissected by multivariate regression confirm uniform distributions of spectral tuning in dLGN and V1. (**A**) Illustration of RRR inputs and outputs. (**right**) Changes in M (green), S (magenta), luminance (black), and hue (blue) contrasts during a single trial period aligned to a neural response. (**left**) Temporal response coefficients (*β*) for each stimulus contrast variable extracted from the same example neurons featured in Figure 3. Insets in the top right of each example plot display the respective sums (*α*) of each *β* for each neuron. (**B**) The collected *β* for all neurons in dLGN and V1 ranked by their |*α*| -hue (**C**) The collected |*α*| for each neuron across stimulus contrast variables ranked by their |*α*| -hue (**D**) Distribution of *r*^2^ coefficients for all neurons recorded. Threshold for significant modulation was set at *r*^2^ = 0.015 (**E**) Maximum mean discrepancy (MMD) for each recording’s *α*M-*α*S distributions extracted by RRR when compared to the distributions generated by our categorical and random models. Median marked by rectangle. Significance measured comparing MMD of experimental distributions to null distribution of MMDs comparing models to themselves (* p¡ 0.05; Wilcoxon rank-sums) (**F**) Aggregated *α*M-*α*S distributions (**above**) with |*α*|-luminance and |*α*|-hue represented as histograms (**below**).

The RRR model identified 2031 neurons (dLGN: 522, V1: 1509) with significant correlation to the stimulus variables (*r*^2^ ¿= 0.015), Figure 4B) and, as with RWA, we observed a broad array of response profiles across both dLGN and V1 (Figure 4Ai-iv, compare to same neurons in Figure 3A,B, Figure 4C). To evaluate each neuron’s overall selectivity for each contrast variable based on RRR, we created a scalar selectivity metric (*α*) by summing the temporal kernels (Figure 4D), though we noted when sorting by the strongest contributions to hue contrast, these neurons also had the strongest contributions to other luminance contrast signaling, in both dLGN and V1 (Figure 4D). As with the RWA, the RRR distributions of M versus S selectivity (Figure 4E) were statistically indistinguishable (Figure 4F) from uniform in both dLGN (H = 0.823, p=1.0) and V1 (H = 0.690, p = 1.0) with no evidence of clustering. The same was true comparing distributions of color versus luminance selectivity (dLGN: H = 0.821, p = 1.0; V1: H = 0.674, p = 1.0).

After confirming similar uniform distributions between the multivariate RRR and univariate RWA, we further evaluated the degree of similarity to our null models. As above for RWA, RRR distributions were scaled before comparison. Comparing the RRR M and S selectivity to the categorical and random selectivity models using MMD, both dLGN (categorical: MMD = 0.011, random: MMD = 0.006) and V1 (categorical: MMD = 0.012, random: MMD = 0.011) distributions showed greater similarity (lower MMD) to the random selectivity model, although the difference in similarity for V1 is notably much smaller (Figure 4E). Comparing the same distributions to those generated with RWA, the dLGN RRR distribution was more similar to its RWA distribution than dLGN (MMD = 0.004 and 0.011, respectively). We found small, but significant positive correlations between tuning for M or S tuning measured by RWA and *α*M and *α*S selectivity estimated by the RRR (M: *ρ* = 0.164, S: *ρ* = 0.232; Spearman’s rank correlation, (Figure 4 with Figure S4A). To compare to the univariate HLSI, we extracted hue and luminance selectivity metrics from RRR fits. We found a very weak, but significant monotonic relationship between —*α*-luminance— (selectivity strength) and HLSI (*ρ* = 0.067, p = 0.003, (Figure 4 with Figure S4B). We observed no relationship with —*α*-hue— whatsoever (*ρ* = −0.020 p = 0.356, (Figure 4 with Figure S4B).

As with the univariate measure, RRR demonstrated the expansion of hue and luminance representations from dLGN to V1. V1 (*α*M: [−1.491,1.713], *α*S: [−1.180,2.069], Figure 4F top) had a broader range of opsin selectivity than dLGN (*α*M: [−0.958,0.811], *α*S: [−0.758,0.821], Figure 4F top) and as a result, a broader range of both hue (dLGN: [−0.496,0.533], V1: [−0.840,0.956] respectively, Figure 4F bottom) and luminance selectivity (dLGN: [−0.858,0.618], V1: [−1.336,1.891], Figure 4F bottom). Contrast selectivity in V1 was strongly right skewed (skew ¿ 1) toward negative values for all types of contrast (mean skew = 1.565 ± 0.531 SD), while dLGN contrast tuning was approximately symmetrical apart from S contrast which was moderately right-skewed (skew = 0.600). Both dLGN and V1 had very strong kurtosis (kurtosis ¿ 5) with V1 (25.591 ± 9.449) having greater kurtosis than dLGN (8.157 ± 1.081) for all contrasts suggesting distributions with sharp central peaks and an abundance of outliers compared to a Gaussian distribution. Interestingly, comparing the distributions of tuning strength (absolute value) for *α*hue and *α*luminance, we found no statistical difference between the distributions in dLGN (D = 0.048, p = 0.588; Komolgorov-Smirnov [KS] Test), but a small but significant difference in the V1 distributions (D = 0.05, p = 0.044). Again, using MMD to compare dLGN and V1 distributions from each recording to the distributions from categorical and random models, we found that the median MMD for both dLGN (Figure 4F; dLGN-random: median disparity = 0.163, p = 8.15E-04; dLGN-categorical: median disparity = 0.050, p = 0.003)and V1 (V1-random: median disparity = 0.148, p = 0.001; V1-categorical: median disparity = 0.123, p = 3.89E-04) indicated greater dissimilarity from the random model, although the experimental distributions all had a significantly greater median MMD than the size-matched null MMDs (p ¡ 0.05, Wilcoxon rank-sums). Overall, we observed significant correlations between the activity of 55% of our recorded neurons (89% recorded during color exchange) and full-field changes in spectral contrast. These correlations reflect a variety of selectivity profiles which, similar to the univariate relationships measured by RWA, were uniformly distributed without evidence of clustering. We observed no preference for luminance or hue tuning in thalamic or cortical populations when directly measured rather than extrapolated from the combination of opsin selectivity, and similar to previous measures (Figure 3) we observed an expansion in contrast selectivity between dLGN and V1.

### Categorical single neuron tuning distribution does not emerge through the cortical hierarchy

In both dlGN and V1, through both univariate (Figure 3) and multivariate (Figure 4) measures, some single neurons showed strong relationships with each form of contrast but with little overall evidence of a categorical population representation of either color or luminance. While this was true of these overall area-based populations, we wanted to ensure that a categorical representation did not emerge within the hierarchy of visual processing within primary visual cortex. It is well known that V1 microcircuits act as intermediate processing steps within the visual hierarchy [39, 40]. As these laminar microcircuits are thought to be responsible for the successive integration of afferent signals, we sought to test if categorical luminance or hue population representations are present within particular laminae even if they do not arise in the total V1 population (Figure 3, Figure 4). The scale of sampling enabled by high-density

### Categorical single neuron tuning distribution does not emerge through the cortical hierarchy

Using normalized entropy, we found that single neuron spectral tuning remains uniform throughout the cortical hierarchy (Figure 5A; RWA-L2/3: H = 0.726, p = 1.0; RWA-L4: H = 0.694, p = 1.0; RWA-L5: H = 0.523, p = 1.0; RWA-L6: H = 0.700, p = 1.0; Figure 5B; RRR-L2/3: H = 0.850, p = 1.0;RRR-L4: H = 0.830, p = 1.0; RRR-L5: H = 0.754, p = 1.0; RRR-L6: H = 0.863, p = 1.0). Using 2-dimensional KS test, we found that only the L6 RWA distribution significantly differed from the aggregated V1 population (Figure 5A; L2/3: D = 0.094, p = 0.150; L4: D= 0.058, p = 0.454; L5: D =0.045, p = 0.581; L6: D = 0.102, p = 0.031), and no RRR distributions differed (Figure 5B; L2/3: D = 0.089, p = 0.220; L4: D= 0.058, p = 0.495; L5: D =0.061, p = 0.240; L6: D = 0.094, p = 0.066). Using HLSI calculated from RWA values, deeper cortical layers had greater skew toward hue contrast preference by the moderate to strong right skew in their selectivity index distributions (Figure 5 with Figure S5; L2/3: 0.694, L4: 1.331, L5: 1.096, L6: 1.892). However, unlike the aggregate V1 population and the RWA distributions, we found no statistical difference in —*α*-hue— or —*α*-luminance— using 1-dimensional KS test (Figure 5B; L2/3: D = 0.058, p = 0.850; L4: D = 0.086, p = 0.128; L5: D = 0.039, p = 0.789; L6: D = 0.085, p = 0.171). Using MMD (as with the collective V1 populations above) to compare each layer to categorical and random models, we found that only L4 and L5 RWA were significantly different from either model (Figure 5C; categorical-L2/3: p = 0.086, random-L2/3: p = 0.147, categorical-L4: p = 0.018, random-L4: p = 0.004, categorical-L5: p = 0.014, random-L5: p = 0.006, categorical-L6: p = 0.090, random-L6: p = 0.122; Wilcoxon rank-sums), with the greater median disparity from the random model indicating more a more categorical-like single neuron representation in L4 and L5. In contrast, RRR distributions across all layers were significantly different from both categorical and random models (Figure 5C; categorical-L2/3: p = 0.002, random-L2/3: p =5.66E-04, categorical-L4: p = 0.001, random-L4: p = 3.89E-04, categorical-L5: p = 8.15E-04, random-L5: p = 5.66E-04, categorical-L6: p = 3.89E-04, random-L6: p = 8.15E-04; Wilcoxon rank-sums)). Although the populations in all layers were neither categorical nor random, RRR also confirmed that L4 and L5, along with the other layers had some bias towards categorical single cell tuning distribution. In total, we do not see evidence for emergence of strictly categorical single neuron representation in any laminar population.

**Figure 5:**
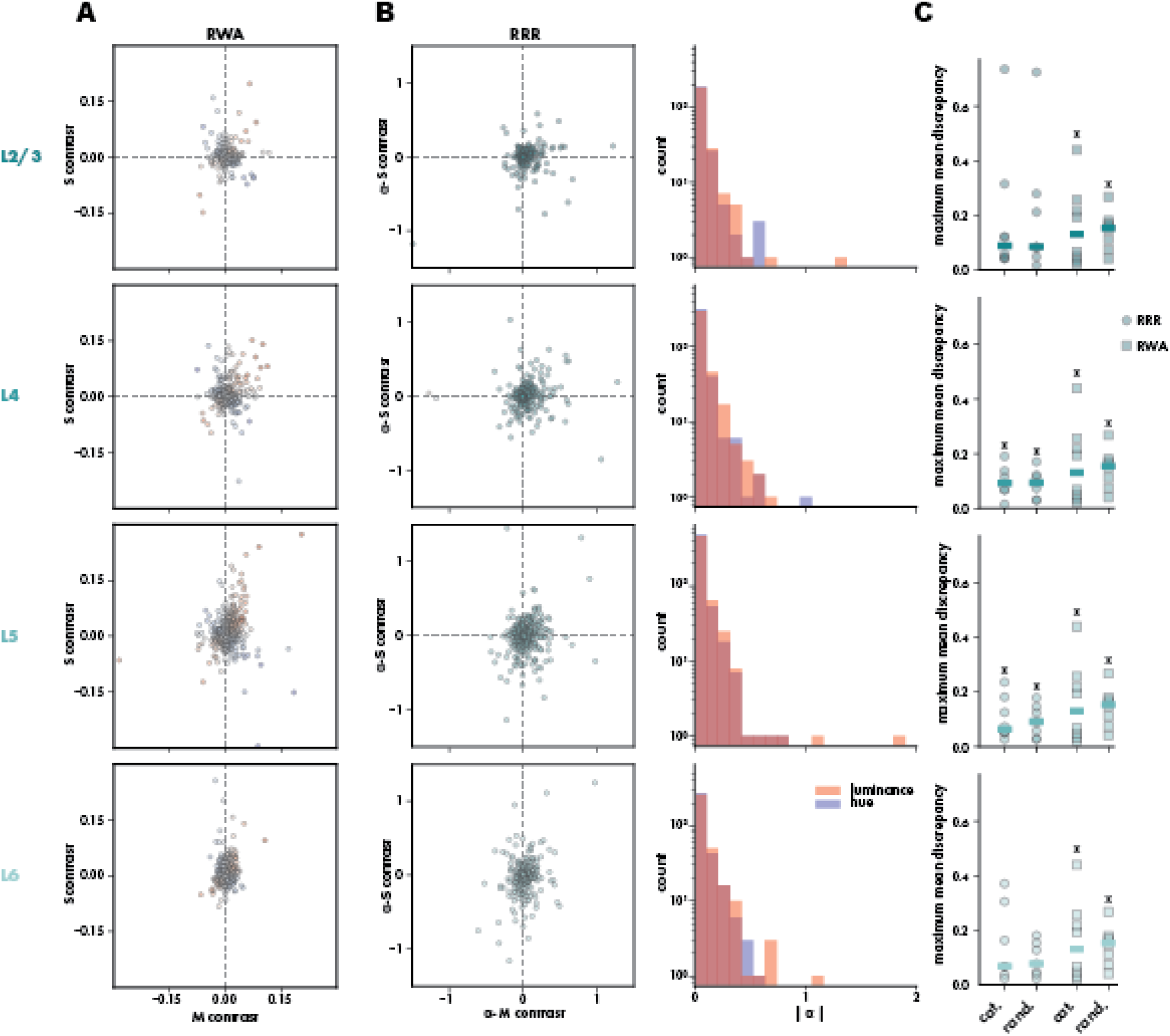
Individual V1 laminae demonstrate similarly uniform distributions of spectral tuning without evidence of functional organization. (**A**) M-S tuning distributions measured using RWA as in Figure 3, broken down by V1 laminae (**B**) *α*M-*α*S distributions extracted using RRR as in Figure 3, broken down by V1 laminae (**C**) MMD comparing the distributions measured in each recording to distributions produced by the categorical and random models. Median marked by rectangle. Significance measured comparing MMD of experimental distributions to null distribution of MMDs comparing models to themselves (* p¡ 0.05; Wilcoxon rank-sums).

### Spectral contrasts can be linearly decoded from populations with uniform tuning distributions

We have thus far shown that population responses in visual thalamus and cortex are comprised primarily of single neurons with mixed selectivity for hue and luminance contrast, and that these neurons form a non-categorical population representation. This raises the question of whether these features could be encoded in a distributed fashion across the population of mixed selective neurons – and if so, how the non-categorical, non-random representation is structured. To first evaluate whether these features are well encoded in the population, we trained support vector machines (SVM) to decode contrast using linear combinations of binned spike data from the various subpopulations in our recordings. Specifically, we used support vector regression (SVR) to decode contrasts as continuous variables rather than discretizing them. All decoders were 5x cross-validated using an 80:20 train-test-split paradigm. To best approximate how well the population is able to encode contrast variables through purely linear combinations of binned spiking activity, we evaluated models using their mean absolute error (MAE) and mean squared error (MSE). While MAE provides a concrete value of how far the model’s estimates are from ground truth, MSE minimizes the effect of small errors and maximizes the effect of large errors to provide a more intuitive metric of decoding. For the purposes of visualization, we reported results as 1 −error (MAE or MSE). Models were evaluated from a minimum of 15 neurons to the maximum population size in steps of 15, at each step in these serial sample additions, populations were resampled 5x to account for single neurons which may have contributed outsized effects to the overall population decoding (Figure 6A).

**Figure 6:**
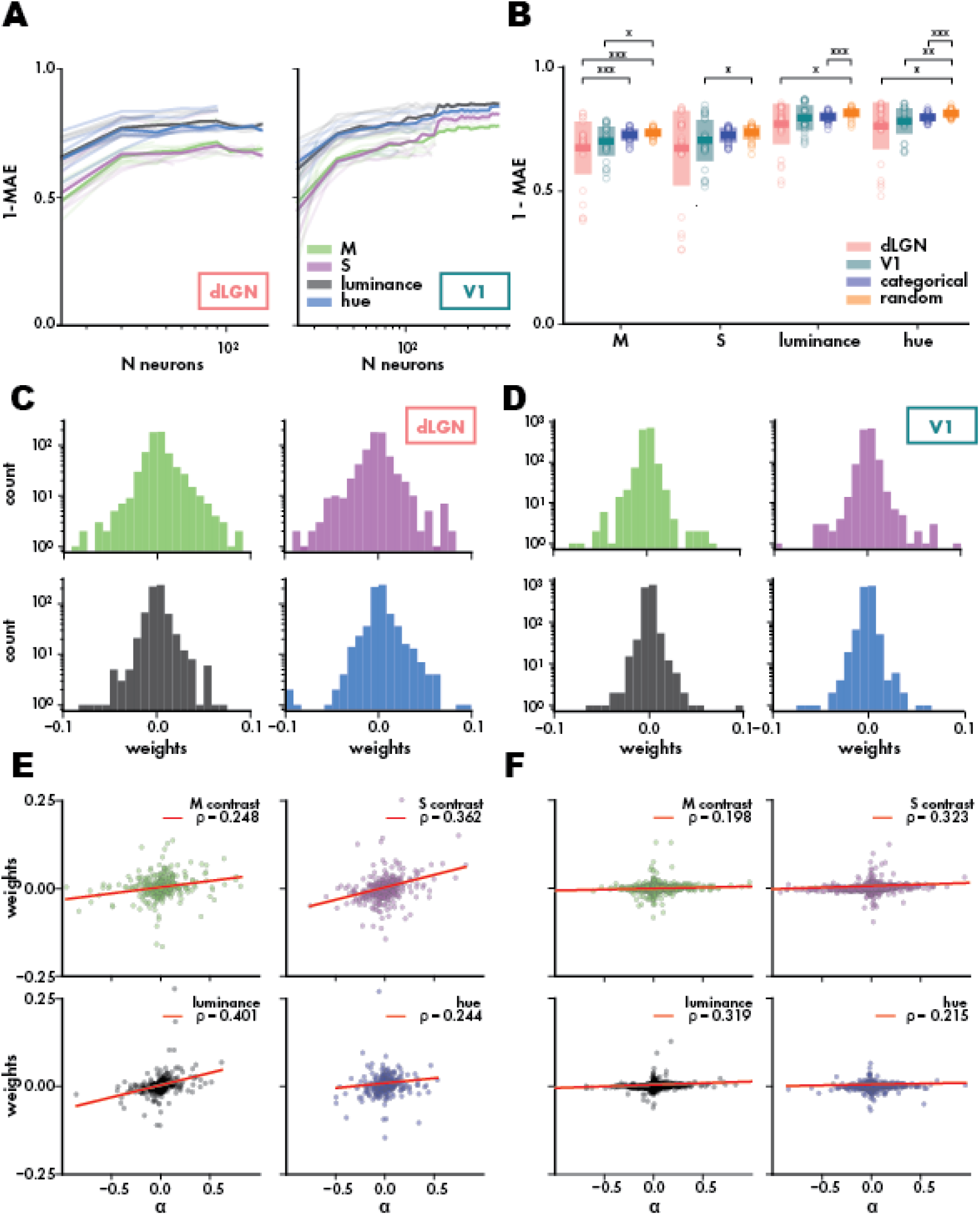
dLGN and V1 populations effectively can represent hue and luminance contrast in relatively small populations of mixed neurons. (**A**) Mean SVR ± performance (1− MAE) for dLGN and V1 as a function of population size. (**B**) Mean ± standard deviation for maximal SVR performance (1 −MAE) across recordings as well as size-matched performances from the categorical and random models. (**E,F**) Distribution of linear decoding weights summed for each neuron at maximal population size for dLGN (**E**) and V1 (**F**) aggregated across recordings. Scatter plots of linear decoding weights for each contrast compared against those neurons’ selectivity measured by RRR (*α*). Each plot is marked with it’s respective spearman correlation (*ρ*) and a linear regression trend line (red).

Decoding performances were compared not only between dLGN and V1, but also against decoding performance from simulated populations from the categorical and random models (Figure 6B). Across contrasts, decoding from both simulated populations was only slightly, but significantly, better than decoding from dLGN (M-contrast decoding: dLGN-categorical p = 7.21E-5, dLGN-random p = 5.49E-7; S-contrast decoding: dLGN-categorical p = 0.189, dLGN-random p = 0.065; luminance decoding: dLGN-categorical p = 0.305, dLGN-random p = 0.002; hue decoding: dLGN-categorical p = 0.007, dLGN-random p = 0.002; 1−MAE, Wilcoxon rank-sums, Bonferroni correction *α*=0.05/6) or V1 (M-contrast decoding: V1-categorical p = 0.039, V1-mixed p = 0.003; S-contrast: V1-categorical p = 0.056, V1-random p = 0.005; luminance decoding: V1-categorical p = 0.305, dLGN-random p = 0.049; hue decoding: V1-categorical p = 0.096, V1-random p = 2.81E-04; 1 −MAE, Wilcoxon rank-sums, Bonferroni correction *α*=0.05/6). We found no significant differences in performance between dLGN and V1.

We investigated the weights learned by each decoder to ask if individual categorical single neurons, or small groups of particularly tuned neurons, had outsized contributions to SVR decoding performance. To specifically look for patterns across neurons, we collapsed the weights for each neuron’s response into a singular value. Weights across regions and contrasts had generally normal distributions (Figure 6C,D), but the Shapiro–Wilk test indicated a statistically significant deviation from normality for all distributions (dLGN-M: p = 6.82E-37, dLGN-S: p = 6.31E-29, dLGN-luminance: p = 3.59E-36, dLGN-hue: p = 2.05E-33; V1-M: p = 6.21E-48, V1-S: p = 2.29E-46, V1-luminance: p = 1.16E-49, V1-hue: p = 2.45E-43). Weights in dLGN were strongly right skewed for all contrasts (skew ¿ 1). Weights in the V1 population were also strongly right-skewed for all contrasts apart from hue which was approximately symmetrical (skew = −0.671). All weight distributions across contrasts and regions had very strong kurtosis (kurtosis ¿ 5) suggesting strong central peaks and an abundance of outlier compared to normal distributions. By comparison, we similarly find that weights for both models, while having approximately normal distributions deviated from normality as defined by Shapiro-Wilk (Figure 6 with Figure S7; categorical-M: p = 1.20E-45, categorical-S: p = 1.24E-44, categorical-luminance: p = 7.38E-52, categorical-hue: p = 1.62E-51; mixed-M: p = 7.75E-46, mixed-S: p = 5.03E-44, mixed-luminance: p = 3.92E-44, mixed-hue: p = 8.64E-45). However, almost all of these distributions were approximately symmetrical (skew ¡ 0.5) with the exception of hue contrast weights in the categorical model (skew = 1.008) and M contrast weights in the randomly selective model (skew = 0.800). All distributions did however, have very high kurtosis (kurtosis ¿ 5).

Elevated kurtosis across experimental samples and models indicates a pronounced concentration of decoding contributions near the mean, much like what was observed for single neuron tuning distributions thus far. While the decoding weights across neurons were relatively symmetrical in the models (with some exceptions), distributions in experimental samples suggest the presence of asymmetric contributions within the population to certain kinds of spectral contrast decoding. In both dLGN (M: *ρ* = 0.248, S: *ρ* = 0.362, luminance: *ρ* = 0.401, hue: *ρ* = 0.244) and V1 (M: *ρ* = 0.198, S: *ρ* = 0.323, luminance:*ρ* = 0.319, hue:*ρ* = 0.215; Spearman correlation), we found that the decoding weights were only weakly to moderately correlated with contrast selectivity. We also found that there was a weak, but significant positive correlation between the strength of neuron weights for luminance and the strength of weights for color (dLGN: *ρ* = 0.342, V1: *ρ* = 0.348), suggesting that at least some of those neurons with greater contributions to decoding luminance also strongly participate in decoding color (Figure 6E,F). The weak correlation between tuning strength and decoding weights the decoders demonstrates that linear decoders are not relying on only the highly-category-selective neurons to achieve accurate performance, and indeed use highly-selective neurons with similar weight to mixed neurons. This suggests that the neural ensemble features driving linear decoding performance may lie in the overall population structure.

### Thalamocortical integration reduces the dimensionality of population representations and changes latent embeddings

The observation of robust population information about luminance and hue – from non-categorical representation formed by mixed contrast selectivity – motivated us to explore the underlying population structure that these decoders were using to extract contrast information. To begin, we quantified the intrinsic dimensionality of neural populations during responses to color exchange stimuli (Figure 7A,B). Intrinsic dimensionality and the geometry of population representations in primary visual cortex have been used as measure of the encoding capacity in primary visual cortex [41], and further used to quantify encoding across modalities, brain regions, and species [38]. High intrinsic dimensionality provide greater computational flexibility for linear readout since the same representation can support a larger number of different output functions.

**Figure 7:**
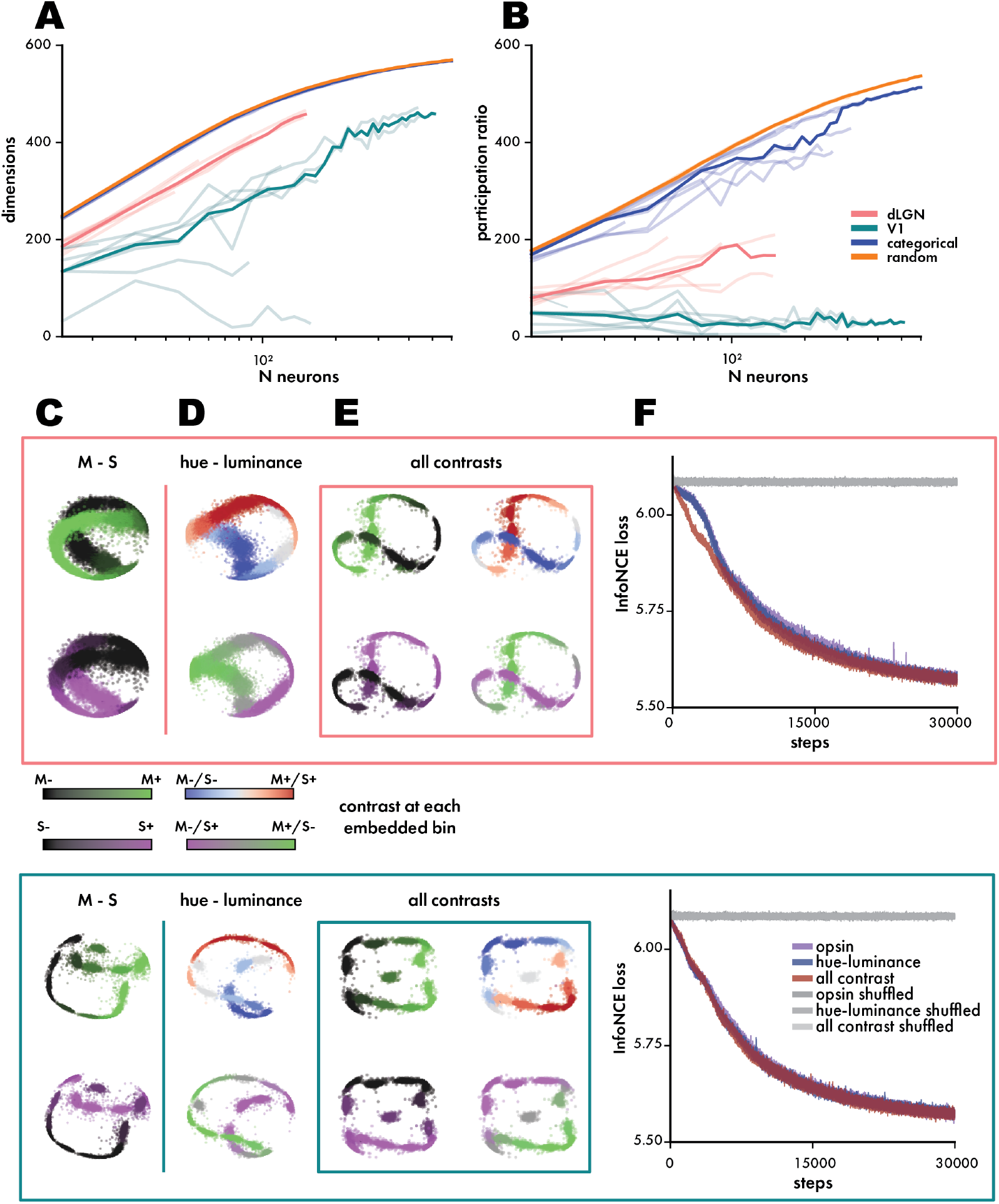
V1 populations show reduced intrinsic dimensionality compared to dLGN or size-matched models. (**A**) Intrinsic dimensionality as measured by PCA as a function of population size. Dimensionality of all recorded populations (lower opacity) overlaid with the mean dimensionality across populations. (**B**) Participation ratio (PR) derived from PCA dimensionality measuring the concentration of variance across dimensions. (**C,D,E**) Latent embeddings of population activity aligned to changes in stimulus variables using CEBRA (dLGN **above**, V1 **below**). Example embeddings from a single recording aligned to changes in M and S contrasts, hue and luminance contrasts, and simultaneous changes to all contrasts (box). Each point in the embeddings represents a trial bin. The embeddings are copied and pseudocolored for each contrast variable put into the algorithm. (**F**) the mean InfoNCE loss across recordings for each embedding.

We measured intrinsic dimensionality first as the minimum number of independent dimensions (eigenvalues) produced by a principal component analysis (PCA) on the covariance of population activity necessary to explain 90% of population variance (Figure 7A). We also expressed PCA dimensionality as a *participation ratio* (*PR*), which measures the sparseness of that population’s dimensionality [42] (Figure 7B). This can also be thought of as *effective dimensionality*, given that dimensionality is weighted by each dimension’s contribution. While a low-dimensional embedding might be indicative of categorical clustering among neurons with similar response profiles, both high and low-dimensional geometries may be compatible with non-categorical responses, if those responses can be mapped onto a low-dimensional manifold in the stimulus space. However, high-dimensional geometries are almost incompatible with categorical responses unless the number of clusters is on the order of the number of conditions. Consistent with the overall trends in decoding performance, both the categorical and random models had higher dimensionality than either experimental population (Figure 7A,B; categorical median = 422, dLGN median = 349.4 [p = 7.16E-10], V1 median = 314.2 [p = 1.92E-21; random median = 426, dLGN median = 336 [p = 4.70E-04], V1 median = 181[p = 7.48E-22];Wilcoxon rank sums with Bonferroni correction, alpha=0.05/6). We found that V1 populations showed statistically lower median PCA dimensionality than dLGN when matched for population size (Figure 7A,B; dLGN median = 336, V1 median = 181, p = 7.16E-10; Wilcoxon rank sums with Bonferroni correction, alpha=0.05/6) with a median difference of 160%. This trend was also seen in the PR of both populations (dLGN median = 119, V1 median = 32.5, p = 1.19E-14; Wilcoxon rank sums with Bonferroni correction, alpha=0.05/6), suggesting that even with lower overall dimensionality, V1 populations are dominated by an even smaller number of dimensions.

While linear dimensionality (as measured by PCA) was high, we considered whether the non-categorical population representation could contain a more structured non-linear low dimensional representations. To do so, we used CEBRA [43], a tool for learning latent embeddings of high-dimensional neural data, to identify low-dimensional nonlinear embeddings related to hue and luminance contrast changes. We found that across subjects, CEBRA was able to extract lower-dimensional embeddings of stimulus contrasts from population activity. We trained CEBRA in supervised manner, using visual stimulus contrasts (M-S, hue-luminance, all contrasts) and embedded the neural activity into 3 latent dimensions. Qualitatively, we find that supervised models of dLGN (Figure 7C,D,E; top) and V1 (Figure 7C,D,E; bottom) produce smooth, structured embeddings with contrast separated along their curves (compare to trial shuffled embedding in Figure 7 with Figure S8). To compare the quality of each non-linear representation (M-S, hue-luminance, and all contrasts) we evaluated the loss of each model (using InfoNCE, see Methods). There was a significant difference in the mean InfoNCE loss between each model and its shuffled counterpart (Figure 7F; dLGN M-S: p = 3.02E-07, dLGN hue-luminance: p = 1.39E-06, dLGN all contrast: p = 3.02E-07; V1 M-S: p = 0.002, V1 hue-luminance: p = 0.002, V1 all contrast: p = 0.002; paired t-test, FDR correction). These embeddings indicate that the population encodes luminance and hue contrast in a non-linear population representation. We found no significant difference in the mean InfoNCE loss between any of the models (Figure 7F; dLGN M-S vs hue-luminance: p = 0.796, dLGN M-S vs all contrast: p = 0.777, dLGN hue-luminance vs all contrast: p = 0.777; V1 M-S vs hue-luminance: p = 0.883, V1 M-S vs all contrast: p = 0.722, V1 hue-luminance vs all contrast: p = 0.661; paired t-test, FDR correction). In summary, we find that single neurons that have mixed selectivity for hue and luminance, form a high dimensional representation of those variables, which can be non-linearly embedded in a separable

## DISCUSSION

In this work we found cone opsin tuning is distributed uniformly among single neurons in the populations of mouse dLGN and V1 and the resulting combinations of cone opsin tuning produce similarly uniform tuning distributions for hue and luminance. Responses to shifts in hue and luminance produce a wide variety of selectivity profiles as defined by multivariate (reduced-rank) regression methods. These decomposed selectivity profiles also further confirmed the uniformity of spectral tuning distributions in the early visual system. Despite tuning across single neurons lacking any clear anatomical or functional organization, populations generate sufficient information to linearly decode stimulus contrasts. Using a self-supervised learning algorithm, we were able to extract consistent low-dimensional embeddings of population activity’s relationship to stimulus contrasts excluding internal or external sources of response variability.

### Mixed selectivity for hue and luminance

Recent work has not only surveyed the distribution of single cell color tuning in the mouse early visual system ([31, 30, 32]), but also the influence of ambient luminance adaptation on the observed distributions [34]. Collectively, these works have suggested a framework of mouse color vision which is able to utilize and transition between a variety of wavelength-sensitive retinal circuit mechanisms which may provide unique ethological advantages [35]. In the process, these studies have also provided evidence of distributed coding of color across many neurons with diverse response types [25, 35, 30]. Here, we quantified the joint coding of hue and luminance as a function of mixed, opsin-specific tuning. While this leaves many questions on the table, it provides the basis for understanding how tuning for specific features might arise from broadly distributed representations and subsequently build complex features such as form and motion.

In our canonical understanding of primate color vision, evidence suggests that color information is encoded and transmitted by distinct cell types beginning at the retina. In this way, color exists at least partially parallel to other visual features driven by changes in luminance [19]. However, there are notable exceptions to this selective wiring for color in primates. Cone selective wiring in primates is a feature of foveal vision, but outside of the fovea, cone signal integration becomes more random as selective wiring becomes impossible. The proportion of strongly hue selective midget cells declines with eccentricity and their sensitivity still tends to be smaller than those found in the fovea [44]. Further, several studies point to mixed single neurons, alongside category specific hue and luminance neurons, in primate V1 [22]. Buzás and colleagues [45] suggest that the overall segregation of cone inputs creating center-surround structure in marmosets was not constituted by local cone cell distributions, but potentially generated by a random wiring model. While a completely random wiring model of the retina isn’t sufficient to explain findings across all studies, it is both consistent with physiological findings across species and potentially a basis mechanism along which other factors introduce the apparent functional segregation seen for cone inputs. In other vertebrate species, color information is encoded by a broader diversity of retinal output channels [7, 15, 46]; many of which are combined with sensitivity toward other visual features. Furthermore, there is mounting evidence of distributed color processing in these species, including mice ([25, 30, 35]). Perceptually, a wide body of evidence points to the co-processing of hue and luminance, including in humans [47–51]. We found neural evidence for co-processing of hue and luminance in V1 and dLGN - single neurons were mixed, but the structure of the population supported separable readout of each feature.

While previously seen as “tolerance” for extraneous variables in otherwise purely selective visual neurons [52, 53], mixed selectivity has been increasingly found as a general property of single cell coding across visual and non-visual brain areas [10, 8]. It has even been suggested that the trial-to-trial variability of neural responses which has previously been discarded as stochastic noise produced as a byproduct of electrophysiology, may itself be reflective of more complex coding strategies which incorporate a broad number of behavioral and perceptual variables, which mutually influence each other’s encoding [54, 12]. Our findings contribute to this growing body of literature which suggests that mixed-selectivity is a fundamental modality of information processing in both cortical and subcortical regions rather than a failure mode of selective single-cell feature coding. Mixed coding may obscure the more straightforward understanding of information encoding at the level of individual and pairs of neurons; it also implies a greater flexibility for information encoding at the level of the larger populations [10, 55]. While our study found that visual populations were sufficiently capable of decoding visual contrasts through the linear combination of binned population activity, we still understand little about the overall strategies, linear or nonlinear, that these populations may be using to encode and decode contrast information in the visual hierarchy.

### Thalamocortical integration of hue and luminance

Our data not only explore the simultaneous representation of luminance and hue at each end of the thalamocortical vision circuit, but also inform other studies of how population codes convey distributed information between regions. This framework of distributed neural population information removes a challenging constraint of finding direct correlations between single neurons and external variables, or between pairs of neurons or cell types. However, it also introduces a level of complexity to our understanding of thalamocortical information processing, which demands large-scale single neuron spiking resolution and will require more advanced computational tools to further disentangle. The differences noted in this study between thalamic and cortical representations of hue and luminance may provide a window into these mechanisms. Aggregating single cell data, we noted a marked expansion in the breadth of the measured tuning distributions in V1 compared to LGN. Prior studies have provided plausible evidence of random or pseudorandom integration of chromatic information in post-retinal processing [25, 45]. Through pseudo-random integration of cone opsin signals, contrast (luminance or hue) information may incidentally arise *de novo*. If this integration process does rely on minimal spatial or cell type bias, a distributed code for hue would be particularly advantageous. While our data do not provide sufficient evidence of a purely random integration of cone opsin signals, the broad, uniform spread of mixed single cell tuning which widens between dLGN and V1 is suggestive of a potentially non-selective integration scheme.

At the population level, we noted a lower intrinsic dimensionality in V1 compared to dLGN during color exchange stimulus. Calculating PR from the measures of PCA dimensionality, we found that the difference between LGN and V1 was even greater, demonstrating that even with lower dimensionality, variance in V1 population activity is dominated by even fewer independent dimensions. While lower dimensional geometries may be indicative of clustered categorical population codes as redundant signals converge, we found no evidence to indicate a clustered organization in the population codes. However, non-clustered representations based on linear mixed selectivity neurons are typically low dimensional and they share similar generalization properties [56]. Different variables are represented by distinct subspaces, allowing for more robust generalization because the coding direction for each variable is independent from the coding for other variables. Low-dimensional, linearly mixed representations have been observed across brain areas [57, 43]. The combined observations of distribution expansion, intrinsic dimensionality reduction, and structured, similarly separable low-dimensional embeddings between dLGN and V1 suggest to us a population coding scheme that is able to integrate spectral information through recombination while preserving information content.

### Limitations of the study

To survey the distribution of color tuning in mouse early visual system, we primarily used a full field color exchange paradigm. There is precedent for the use of full field stimuli for surveying color pathways [30] and there are clear advantages to this stimulus including its simplicity of implementation, the ability to simultaneously survey color tuning across retinotopic space, and the relative simplicity of analysis without the confounding variable of spatial structure. However, the limitations of this stimulus are implicit in its advantages. A major component of signal integration in the visual hierarchy is spatial integration. Across species, many mechanisms of chromatic signal integration rely on comparisons of wavelength selectivity between the center and surround of each cell’s receptive field [58, 35]. In the case of strongly antagonistic center-surround receptive field structures, full-field contrast changes might cancel out otherwise robust responses. As well, many neurons in V1 are known to be more strongly, or exclusively, driven by spatial structure, although prior work exploring color tuning in mouse V1 specifically, has noted that spatial frequency in color opponent neurons is lowpass [25]. Nonetheless,our experimental paradigm reliably elicited hue and luminance dependent responses from single neurons, demonstrating a broad, uniform distribution of single cell tuning that is consistent with distributions surveyed in studies with spatially structured stimuli [35].

## METHODS

### Animals

All animal procedures were approved by the University of Colorado Anschutz Medical Campus Institutional Animal Care and Use Committee (IACUC) according to ARRIVE 2.0 guidelines. Mice (n=8) in this study were male C57BL/6J aged 60 – 182 days. For all mice, an initial headpost procedure was performed to attach a 10mm diameter circular opening aluminum head-fixation plate to the skull above the left hemisphere.

### Spiking Models

To generate statistical expectations for the selectivity profiles of purely selective or randomly selective single neuron opsin tuning distributions, we constructed populations of simulated neurons with fixed profile Gaussian tuning in a two-dimensional contrast space. Each neuron’s response was defined by it’s preferred contrast vector, [*M, S*] and a tuning width parameter controlling sensitivity to deviations from the preferred contrast. Given a stimulus contrast vector, the instantaneous firing rate of each neuron was computed using a Gaussian tuning function:

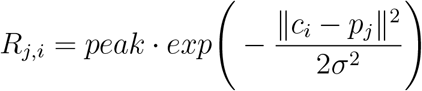

where *R*_*j,i*_ denotes the response magnitude of neuron *j* to stimulus contrast *c*_*i*_, and *p*_*j*_ is the neurons preferred contrast. The parameter *σ* controls the tuning width, and *peak* is the maximum evoked firing rate.

Neural responses were modeled as *homogeneous Poisson processes* with a constant firing rate during the stimulus window. While this is less analogous to biological neuron responses, this assured our null models would not introduce variables of spiking dynamics or varied response profiles while still reliably producing response metrics based on trial summation and averaging (see Response Weighted Average below). The total trial duration was divided into three segments: a pre-stimulus baseline period, a stimulus presentation window, and a post-stimulus period. Each bin was 10ms in duration. Spikes during the pre- and post-stimulus windows were generated using a fixed baseline firing rate, while spikes during the stimulus window were generated with the sum of baseline and evoked rates:

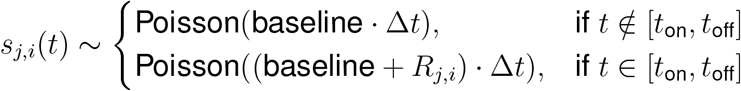

Because the evoked rate was constant during the stimulus window, this model generated spike trains with *flat, box-like PSTHs*, lacking the smooth temporal structure typically observed in real cortical recordings.

### Electrophysiological recordings

On the day of electrophysiological recording, the animal was anesthetized and placed in a stereotaxic. Burr hole craniotomies were made over primary visual cortex (V1) and higher visual cortical areas for those probes targeting lateral geniculate nucleus (LGN). V1 burr holes were centered on stereotaxic coordinates −3.1 mm anterior-posterior (AP), −2.9 mm medial-lateral over the left hemisphere visual cortex. For visual experiments targeting LGN, the electrode was mounted at 10-15° tilt AP and the target stereotaxic coordinates were −2.0 mm, −2.5 mm (AP,ML). The animal was allowed to recover on the electrophysiology rig during Neuropixels probe insertion. Neuropixels probe insertion was done using feedback-controlled piezoelectrical manipulators (New Scale Technologies). After breaking the surface, Neuropixels were lowered to 1.2 mm (V1 recordings only), 3.5mm (electrical stimulation and thalamic targeting), and 800 *µm* (V1 Neuropixels 2.0 recordings). Neuropixels data were acquired with the Open Ephys GUI at 30kHz (AP band and Neuropixels 2.0 wide band) and 2.5kHz (LFP band). Stimulus timing was synchronized using a multi-function data acquisition card (National Instruments) through Open Ephys [59]. All data were processed through Kilosort 2.5 [60] to isolate putative single neurons from Neuropixels recordings. Individual experiments were packages into Neurodata Without Borders (NWB, [61]) files containing sorted unit spike times and stimulus time information.

#### Estimated anatomical labels

Trajectory planning for each experiment was done using Pinpoint and probe positions were adjusted to match the planned trajectories. In each recording, we used two Neuropixels 1.0 probes which descended through the cortex down to dLGN. We additionally used a 4-shank Neuropixels 2.0 probe which was only inserted into V1 (Figure 2 with Figure S1A). Angles of approach and insertion remained the same across recordings although there was some variance in the exact stereotaxic location of insertion. Local field potentials (LFP) were decomposed into their power spectra and the transition from brain to air was determined by locating a sudden drop in LFP power (Figure 2 with Figure S1B). Multiunit amplitudes (MUA) from spiking activity were used to identify peaks which could be used to localize anatomical structures. Depending on the trajectory and depth of insertion, we would regularly see a very large peak at CA1 of the hippocampus and a smaller peak at L5 of the visual cortex. These landmarks along with the surface were used to align and correct estimations of channel position (Figure 2 with Figure S1C). Given entry point, tip point, trajectory angles, and electrophysiological landmarks, we were able to model the relative position of all probe channels in 3D space. Channels were each assigned to an anatomical structure according to the Allen Common Coordinate Framework [62] (CCF) (Figure 2 with Figure S1D).

#### Sorting quality Metrics

Quality metrics were calculated for all putative neurons using Bombcell [63] and filtered from noise and multi unit activity using previously validated thresholds (Figure 2 with Figure S2). Units were first classified as noise or not noise based on thresholds measures of each unit’s waveform including peaks and troughs, duration, baseline flatness, and spatial decay slope (Figure 2 with Figure S2A). Units were then classified as likely being multiunit amplitudes (MUA) or good, clean neurons based on each unit’s spiking behaviors (Figure 2 with Figure S2B).

### Visual Stimuli

Visual stimuli were presented across the full visual field using a custom spherical stimulus enclosure. A custom DLP-projector designed for the mouse stimulus system [31] provided independent spatiotemporal modulation of ultraviolet and green light. The projection system operated at 1024 x 768 pixel resolution and a refresh rate of 60Hz, achieving a maximum intensity in the high-mesopic (400 photons(*γ*)/s) to photopic (15000 *γ*/s) range (green-M: [292.4*γ*/s, 14328 *γ*/s, 30596.05*γ*/s], green-S: [774.93*γ*/s, 1610.49 *γ*/s, 2485.06*γ*/s], UV-M: [420.21 *γ*/s, 3257.00 *γ*/s, 5449.66 *γ*/s], UV-S: [1105.42 *γ*/s, 8207.82 *γ*/s, 13901.64 *γ*/s]; [min, 50%, max]).Planar stimuli were spatially warped according to a custom fisheye warp for spherical projection. The fisheye warp was created through an iterative mapping protocol using the meshmapper utility (http://paulbourke.net/dome/meshmapper/). Custom stimuli were generated using PsychoPy [64].

#### Calibrating stimuli for opsin specific stimulation

We used a custom projector with green and UV LEDs whose spectra overlapped with the sensitivity spectra of mouse M and S opsin (Figure 2 with Figure S1A). Using methods outlined in previous work [36], we used an Apogee Instruments PS-100 spectroradiometer to measure the intensity of each LED from the vantage point of the headfixed mouse in our stimulus environment. We then modeled the relative stimulation of M and S opsin at each step of LED power within their 8-bit ranges. Importantly, we also measured the effect of green LED on S opsin and vice versa to account for spectral overlap (Figure 2 with Figure S1B). Using the modeled opsin stimulation data, we calculated the relative M-S contrast at each combination of green-UV LED intensities and mapped out the potential stimuli within M-S contrast space:

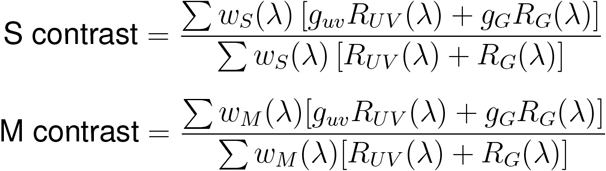

*w*_*S*_(*λ*), *w*_*M*_ (*λ*): The S- and M-opsin sensitivity functions[65] *R*_*UV*_ (*λ*), *R*_*G*_(*λ*): Radiance of UV and green LEDs *g*_*uv*_, *g*_*g*_: contrast of UV and green LEDs

For each experiment, we selected radially symmetrical contrasts within that potential stimulus space so as to not bias our measurements in any particular tuning direction (Figure 2 with Figure S1C).

#### Luminance Flash

The luminance flash stimulus was spatially uniform covering the full visual field. Flashes alternated between 100% and 0% luminance contrast. Between flashes, the stimulus maintained 50% luminance contrast. There were 100 flashes in each stimulus (50 bright, 50 dark), with a 500ms flash duration and 2.5s between flashes.

#### Full-Field Color Exchange

The full field color exchange presented a uniform color across the full visual field which changed randomly every 500ms. Colors were drawn from a pre-defined distribution which was radially symmetrical in M-S contrast space (Figure 3 with Figure S3). In each stimulus, there were between 680-3400 presentations depending on the mouse’s tolerance.

### Data Analysis

All data analysis was performed in Jupyter notebooks (python 3.12). Each data analysis notebook is housed in a github repository (https://github.com/denmanlab/mixedcolor).

#### Color Tuning Measurements

To estimate each neuron’s preferred color direction in M-S opsin space, we used response-weighted averaging RWA [37], described mathematically as:

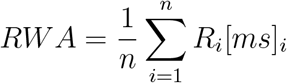

*n* is the total number of tested stimuli, [*ms*]_*i*_ is the opsin contrasts of the *i*^*th*^ stimulus, and *R*_*i*_ is the response to the *i*^*th*^ stimulus. The response is defined as the z-scored (from baseline) mean firing rate, divided by the sum of all responses, normalizing responses to sum to 1:

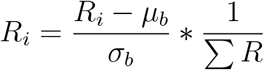

To account for potential inconsistencies in the radial symmetry of presented contrasts, we multiplied [*ms*]_*i*_ with a whitening matrix, and then transformed the RWA back to the original color space.

The RWA underestimates the true preference strength of neurons. This is the result of a number of factors including the width of a neuron’s tuning across a given contrast spectrum, the stimulus’ coverage across the full feature space (and the symmetry of that coverage), spiking variability, and normalization inherent in the RWA formula. This was confirmed with a spiking model (see above) which generated Poisson spike trains with pre-defined preferences to stimulus contrasts. Even neurons with selectivity for maximal opsin contrasts still held their RWAs within a range of [−0.2,0.2] when measured against our stimuli.

#### Hue-luminance Selectivity Index

The color-luminance selectivity index (HLSI) measures how well each point in M-S contrast space is aligned to either the color or luminance axes. The color and luminance (l) axes are rotated 45 degrees from the original M-S basis, so to adjust the coordinates to color-luminance space:

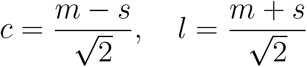

To calculate the HLSI:

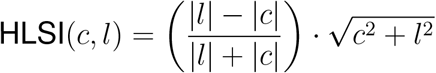

#### Reduced-Rank Regression Linear Encoding Model

We implemented the reduced-rank regression model used by Posani et al., 2025 [38]. Code for implementing and fitting this model can be found at https://github.com/realwsq/brainwide-RRR-encoding-model.

For each neuron, *n*, the RRR describes it’s temporal responses as a linear, time-dependent combination of input variables. The extracted temporal bases for each contrast 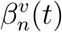 were summed to measure the total selectivity (*α*) of neuron *n* to contrast *v*.

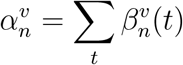

Sharing these temporal bases across neurons and input variables significantly reduces the number of regression parameters. In cases where the goal was to measure the absolute modulation of a contrast, we used the absolute sum of the time-varying coefficient.

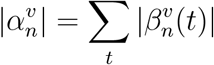

Hyperparameters for each model fit were selected iteratively (500 iterations) with a 70:30 train-test split.

#### Maximum Mean Discrepancy

To quantify differences between multivariate distributions, we used *maximum mean discrepancy* (MMD) — a non-parametric, kernel based statistic that measures the distance between two probability distributions based on their samples. Specifically, MMD assesses how *dissimilar* two sets of observations are by comparing their mean embeddings in a reproducing kernel Hilbert space (RKHS). The MMD is zero if the two distributions are identical. We used the radial basis function (RBF) kernel to compute MMD, which is sensitive to both local and global differences in distributional structure. The kernel bandwidth was set using the *median heuristic*, defined as the inverse of twice the median of the pairwise Euclidean distances among pooled samples from both distributions. To account for differences in scale across features, all data were normalized to have zero mean with unit variance. To assess the statistical difference between the distributions produced by spiking data versus models, we compared the observed MMD to size matched null MMDs (model distributions compared against themselves) using the Wilcoxon rank-sum test for significance.

#### Linear Stimulus Decoding from Population Activity

To linearly decode stimulus contrasts from population activity, we used support vector regression (SVR) models implemented using the scikit-learn Python package. For input, the SVR takes a 1-dimensional features vector (*y*) that consists of the change in contrast for each trial (*contrast*_*n*_ −*contrast*_*n−*1_). Vector *y* is compared to a samples vector (*X*) with shape trials x neurons*bins. For this study, we used a 10ms bin size for all SVR decoding tasks. Each recording and model was evaluated individually rather than pooling populations. For evaluations of performance versus population size, samples from each model were sized matched to experimental populations.

#### Measures of Dimensionality

Intrinsic dimensionality was measured by decomposing the co-variance matrix of binned population activity across trials using principal component analysis (PCA). The *participation ratio* (PR) quantified the effective dimensionality of data by measuring how evenly the variance is distributed across the eigenvalues, *λ*_*j*_ (i.e. dimensions), of the PCA decomposition:

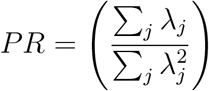

where *λ*_*j*_ is the j-th eigenvalue fo the covariance matrix ranked by the amount of variance explained. A higher PR indicates that variance is more evenly spread across multiple dimensions, suggesting a higher effective dimensionality of the population representation.

#### Latent embeddings of contrasts in population activity

Low-dimensional embeddings of population activity were generated with CEBRA [43] using their PyTorch API. CEBRA is a self-supervised learning algorithm that provides consistent, interpretable low-dimensional embeddings of high-dimensional neural population data aligned to temporally labeled stimulus variables. Using InfoNCE (noise contrastive estimation) loss, CEBRA aims to classify correct positive samples from multiple negative samples. Data for all recordings was trial matched to the minimum number of trials in an experiment (680 trials). Spiking data and stimulus data were binned at 10ms. One out of eight recordings was excluded from analysis for poor model convergence caused by small population samples (n = 7). Population activity from all regions and recordings were embedded using the same model architecture and parameters. Data loading and model parameters are included in Table 1.

**Table 1:** CEBRA parameters. Data loading and model parameters used in CEBRA runs.

| Data Loading Parameter | Value |
| --- | --- |
| *num_steps* | 30000 |
| *batch_size* | 512 |
| *conditional* | 'time_delta' |
| *time_offset* | 10 |
| Model Parameter | Value |
| *model_architecture* | 'offset10-model' |
| *distance* | 'cosine' |
| *num_hidden_units* | 128 |
| *output_dimension* | 3 |
| *temperature* | 1 |
| *learning_rate* | $3 \times 10^{-4}$ |

## RESOURCE AVAILABILITY

### Lead contact

Requests for further information and resources should be directed to and will be fulfilled by the lead contact, Daniel J. Denman.

### Materials availability

This study did not generate new materials.

### Data and code availability

All original code has been deposited on GitHub (https://github.com/denmanlab/mixedcolor).

## ACKNOWLEDGMENTS

This work was funded by R00EY028612 (DJD) and F30EY034775 (JSM). The authors thank all members of the lab for their support.

## AUTHOR CONTRIBUTIONS

Conceptualization, J.S and D.J.D.; methodology, J.S and D.J.D.; investigation, J.S and D.J.D.; writing-–original draft, J.S and D.J.D.; writing-–review & editing, J.S N.G., and D.J.D.; funding acquisition, J.S and D.J.D.; resources, D.J.D; supervision, D.J.D.

## DECLARATION OF GENERATIVE AI AND AI-ASSISTED TECHNOLOGIES

During the preparation of this work, the author(s) used ChatGPT adn ClaudeAI in order to streamline writing and debugging analysis code. After using this tool or service, the author(s) reviewed and edited the content as needed and take(s) full responsibility for the content of the publication.

## SUPPLEMENTAL INFORMATION INDEX

Figures S1-S12 and their legends.

## SUPPLEMENTAL FIGURES AND LEGENDS

**Figure S1:**
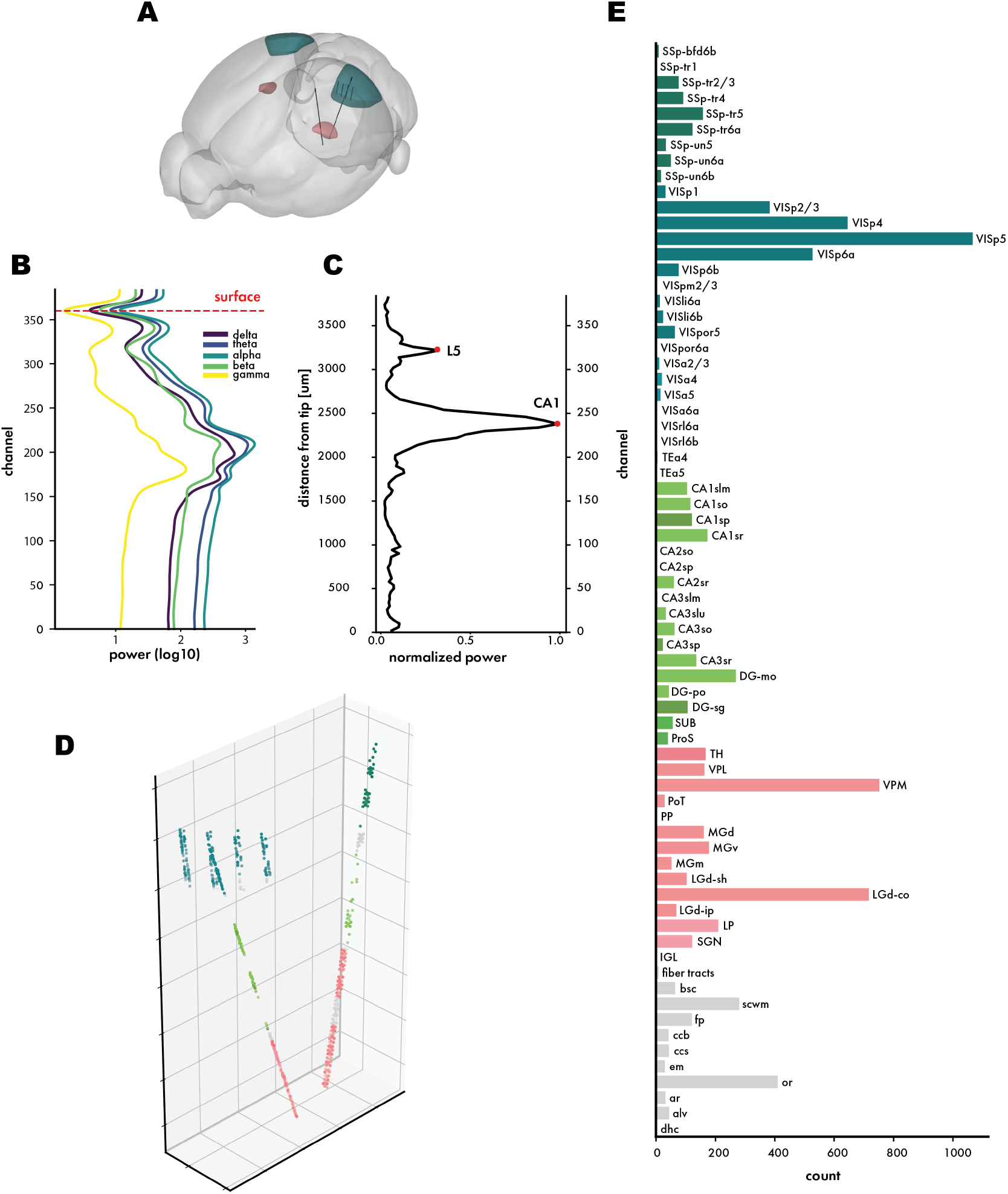
Estimated anatomical labeling of neurons using electrophysiological landmarks. (**A**) Illustration of probe trajectories for a single recording. (**B**) Local field potential (LFP) power spectra from an example recording decomposed into spectral bands and used to approximate the channel where the probe transitions from brain to saline via power minimum near the putative surface. (**C**) Multiunit amplitudes (MUA) from spiking activity used to identify peaks (red dots); peaks were used to localize neocortical layer 5 and the CA1 pyramidal cell layer. (**D**) The relative position of channels from all three probes in the example recording shown in **A** in 3D space. Channels are colored by their estimated anatomical structure according to the Allen Common Coordinate Framework (CCF). (**E**) Bar plot of all neurons recorded across 8 recordings grouped by their estimated anatomical region. Regions are roughly ordered by relative depth.

**Figure S2:**
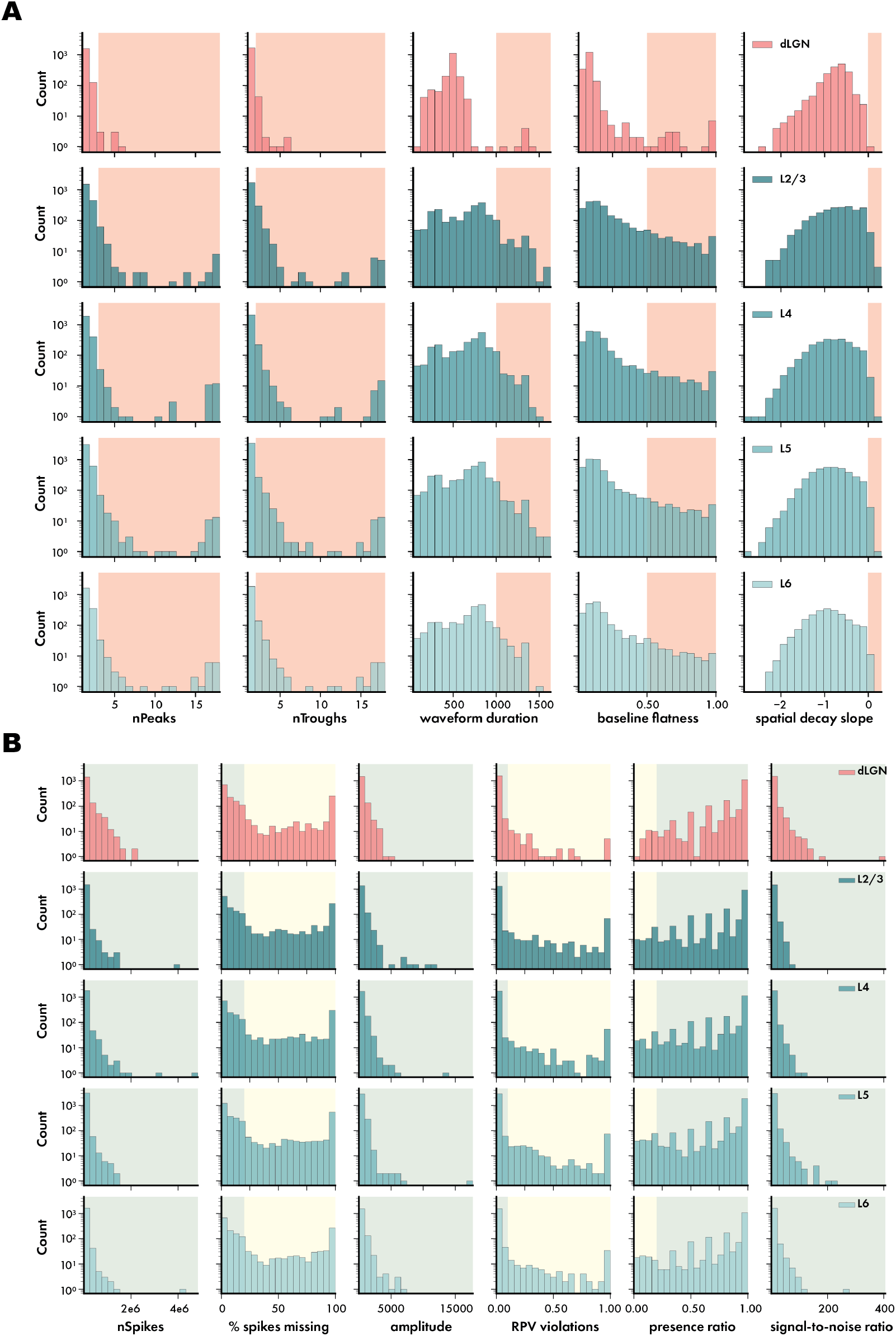
Automated spike sort curation using Bombcell. (**A**) Distribution of waveform morphology metrics across units in dLGN and each layer of V1 classified as noise (shaded red) or not noise based on thresholds measures of each unit’s waveform including peaks and troughs, duration, baseline flatness, and spatial decay slope. (**B**) Units classified as multiunit amplitudes (MUA, shaded yellow) or good, clean neurons (shaded green) based on spiking behaviors.

**Figure S3:**
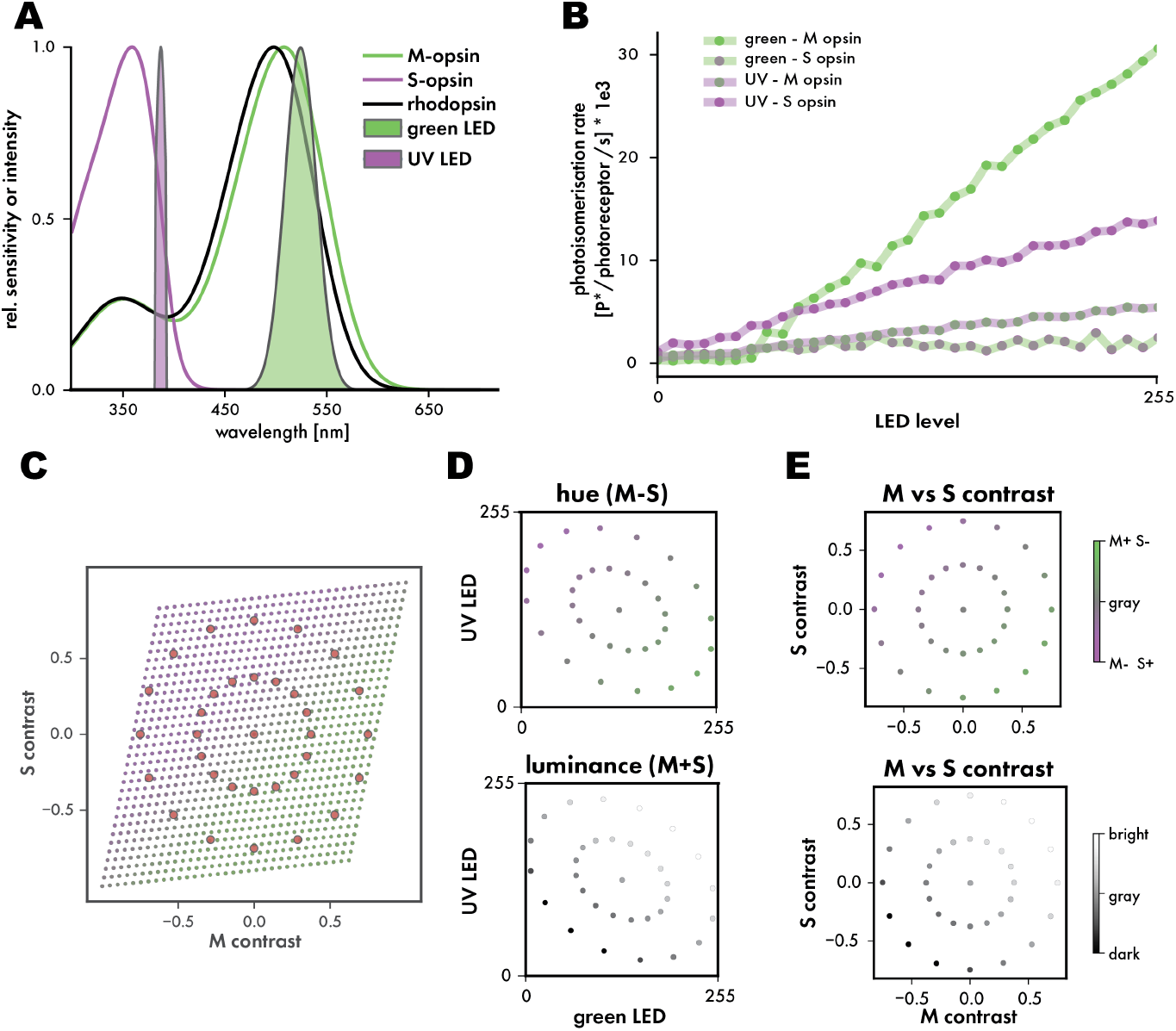
Opsin-specific stimulus calibration. (**A**) Mouse cone opsin spectra [65] overlaid with emission spectra of green and UV projector LEDs. (**B**) Modeled measurements of M and S opsin stimulation (photoisomerization rate) based on spectroradiometer measurements of LED intensity from the vantage point of the headfixed mouse in our stimulus environment. (**C**) The space of potential M-S opsin contrast based on the range of LED intensities produced by our projector. Overlaid are example points for radially symmetrical stimulus contrast parameters (red). (**D,E**) The same example stimulus parameters from **C**, projected into LED intensity space and the derived M-S contrast space colored by their relative hue contrast (green, magenta) or luminance contrast (black, white).

**Figure S4:**
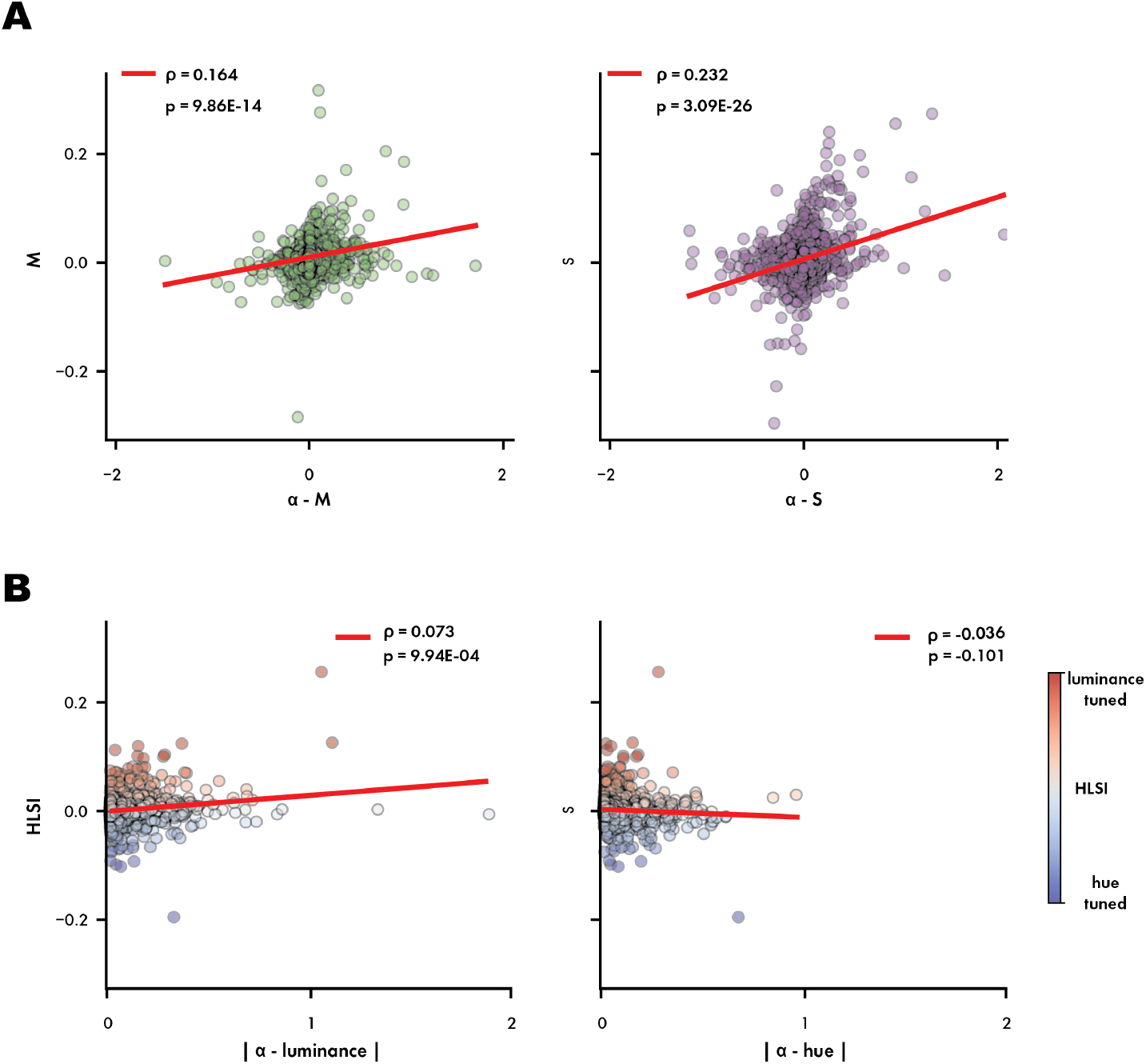
Correlations between response weighted average (RWA) and reduced rank regression (RRR) selectivity metrics. Correlations between response weighted average (RWA) and reduced rank regression (RRR) selectivity metrics. (**A**) Scatter plots of *α*M and *α*S against those neurons’ selectivity measured by RWA. Each plot is marked with it’s respective Spearman correlation (*ρ*), associated p value, and a linear regression trendline (red) (**B**) Scatter plots of *α*-hue and *α*-luminance against those same neurons HLSI calculated from their RWA M-S tuning. Spearman correlation, p value, and trend line marked as in A. Points colored by their respective HLSI value (color bar).

**Figure S5:**
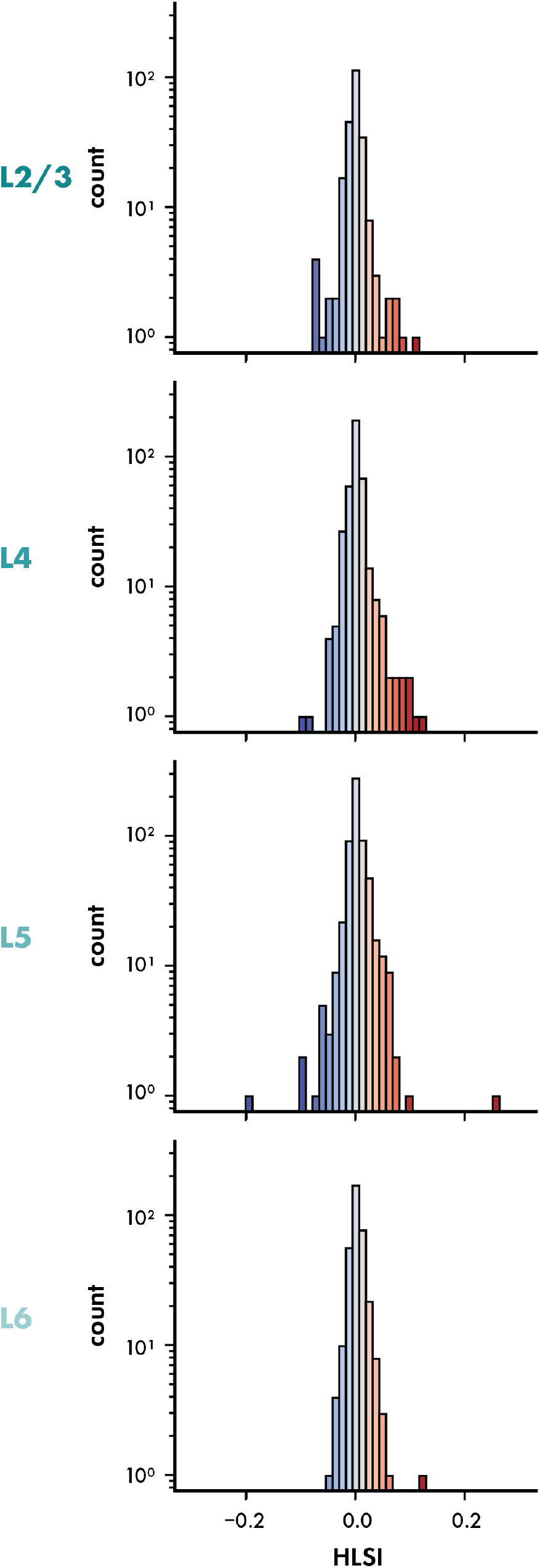
HLSI across layers based on RWA. Histograms of HLSI for each cortical layer, colored by HLSI value as in Figure 3C.

**Figure S6:**
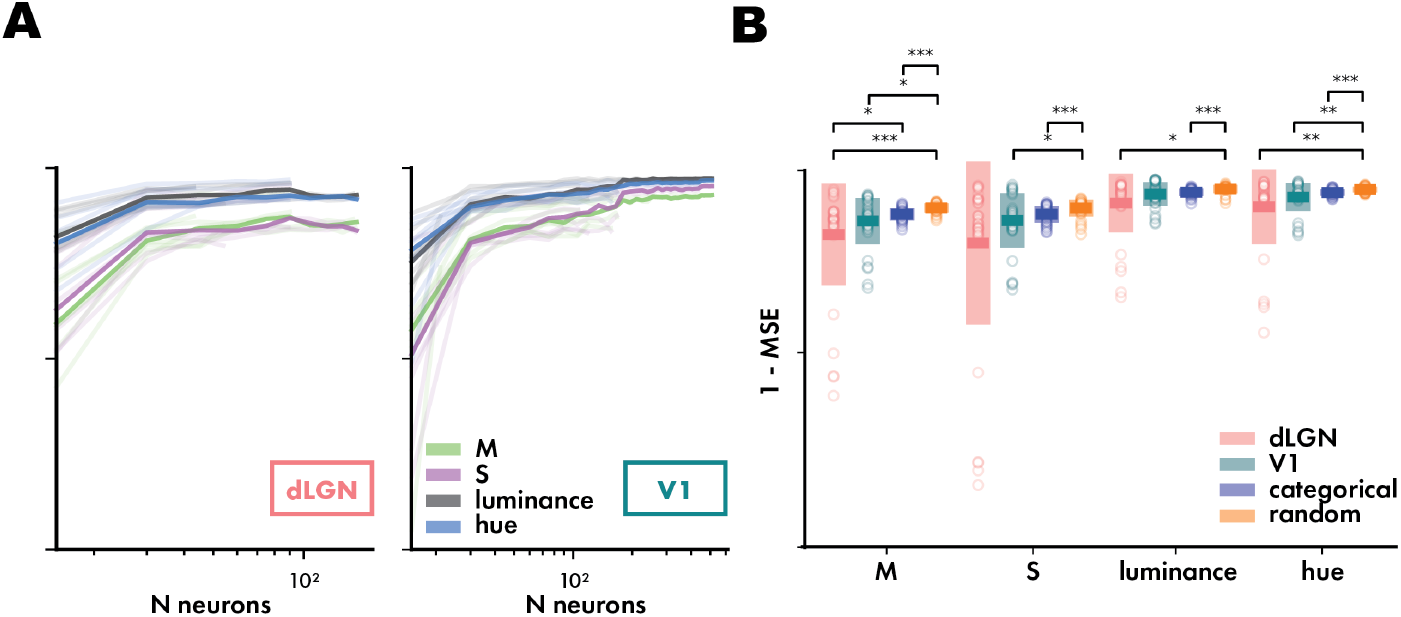
SVR decoding performance measured using mean squared error (MSE). (**A**) Mean SVR performance (1− MSE) for dLGN and V1 as a function of population size. (**B**) Mean ± standard deviation for maximal SVR performance (1− MSE) across recordings as well as size-matched performances from the purely selective and mixed selective models. (* p¡ 0.05; Wilcoxon rank-sums).

**Figure S7:**
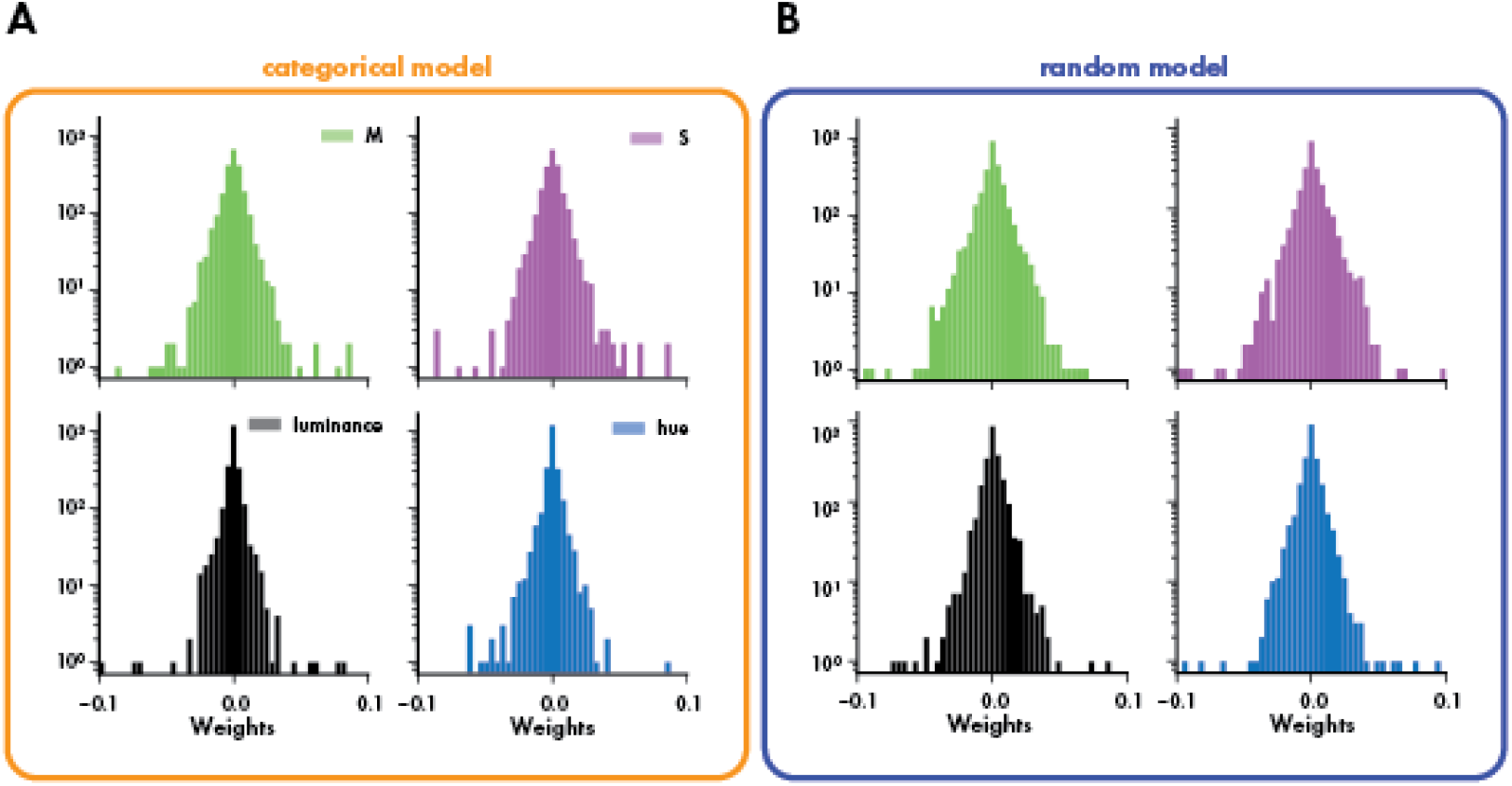
Figure 6 - Supplement 2. Null model linear decoding weights. Distribution of aggregated categorical (**A**) and random (**B**) model weights for contrast decoding (as in Figure 6C,D).

**Figure S8:**
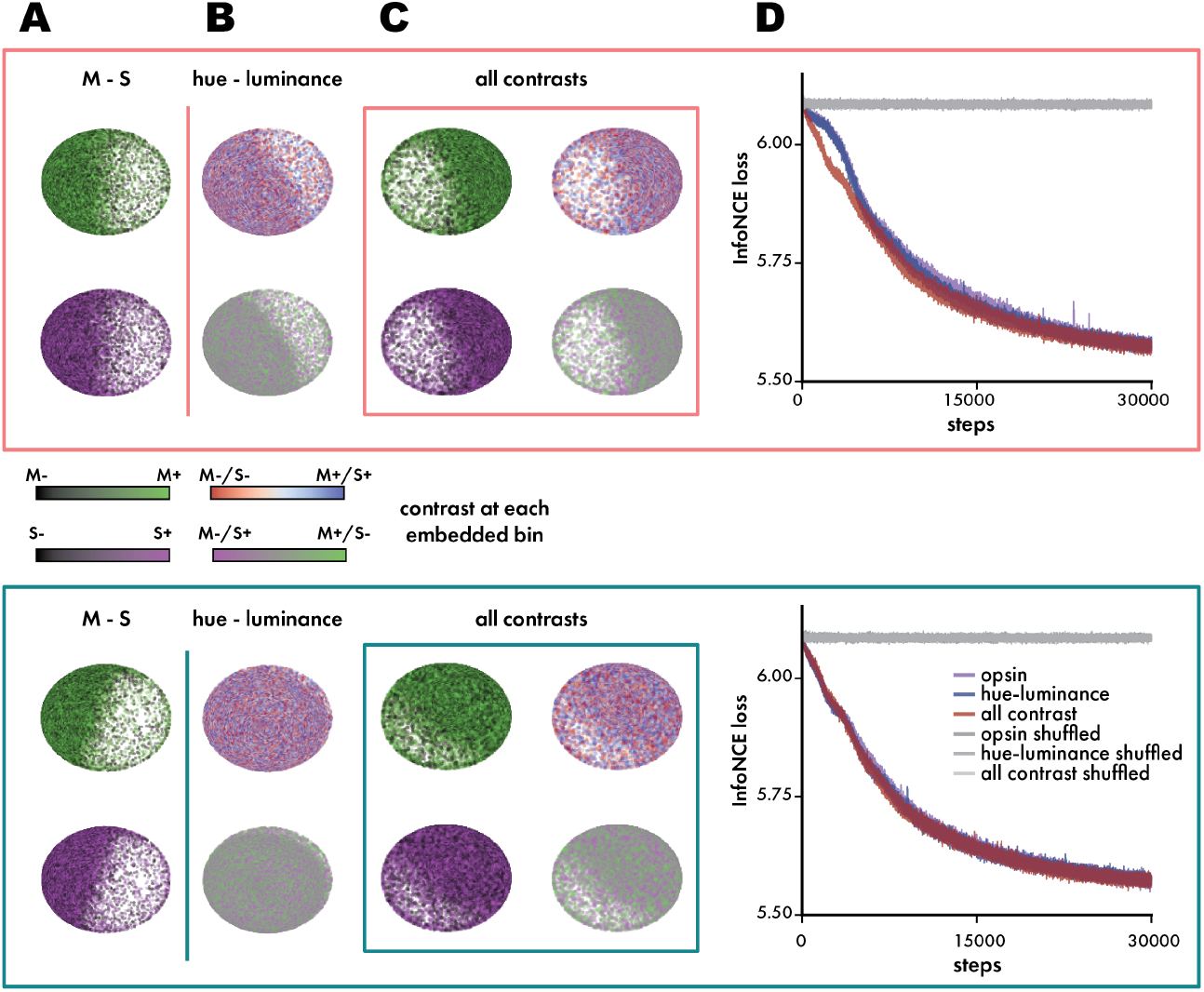
Latent embeddings of population activity fit with shuffled trial structure. (**A,B,C**) As in Figure 7C, we plotted the latent embeddings from the same sample recording which were fit using shuffled trial data. (**D**) The InfoNCE loss curves for all embeddings as plotted in Figure 7F. Curves for shuffled models plotted in grey.

Please define all scale and error bars, and please review the Cell Press figure guidelines before submission: https://www.cell.com/figureguidelines. Example figure created by Cassie Comeau, Cell Press.

**Figure S9:**
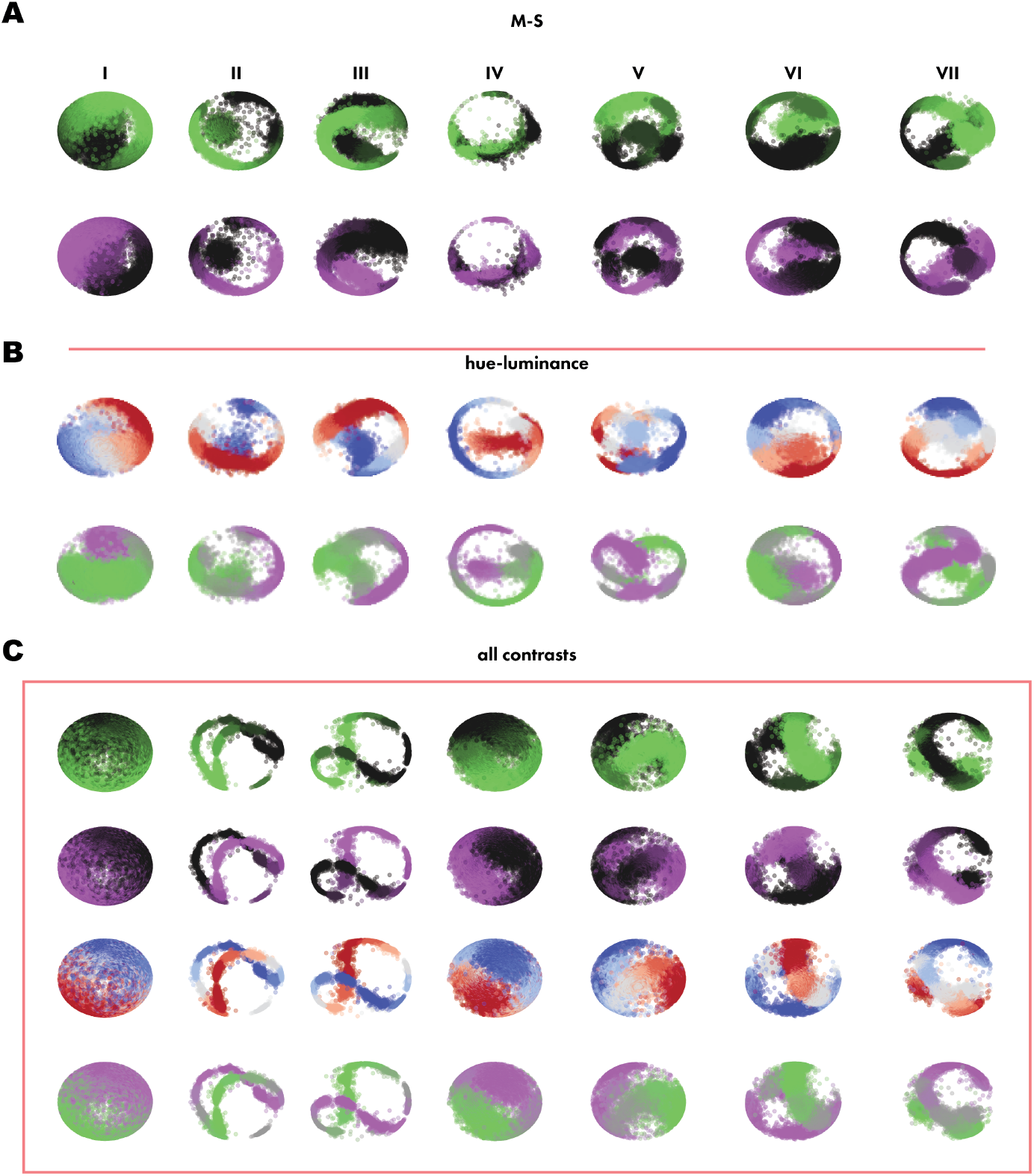
Latent embeddings of dLGN population activity for all recordings. Plots of dLGN population activity from all recordings (n = 7, I - VII) embedded against. (**A**) M-S contrasts, (**B**) hue-luminance contrasts, (**C**) all contrasts. Recording III was used as the example for Figure 7 and Figure 7 with Figure S8.

**Figure S10:**
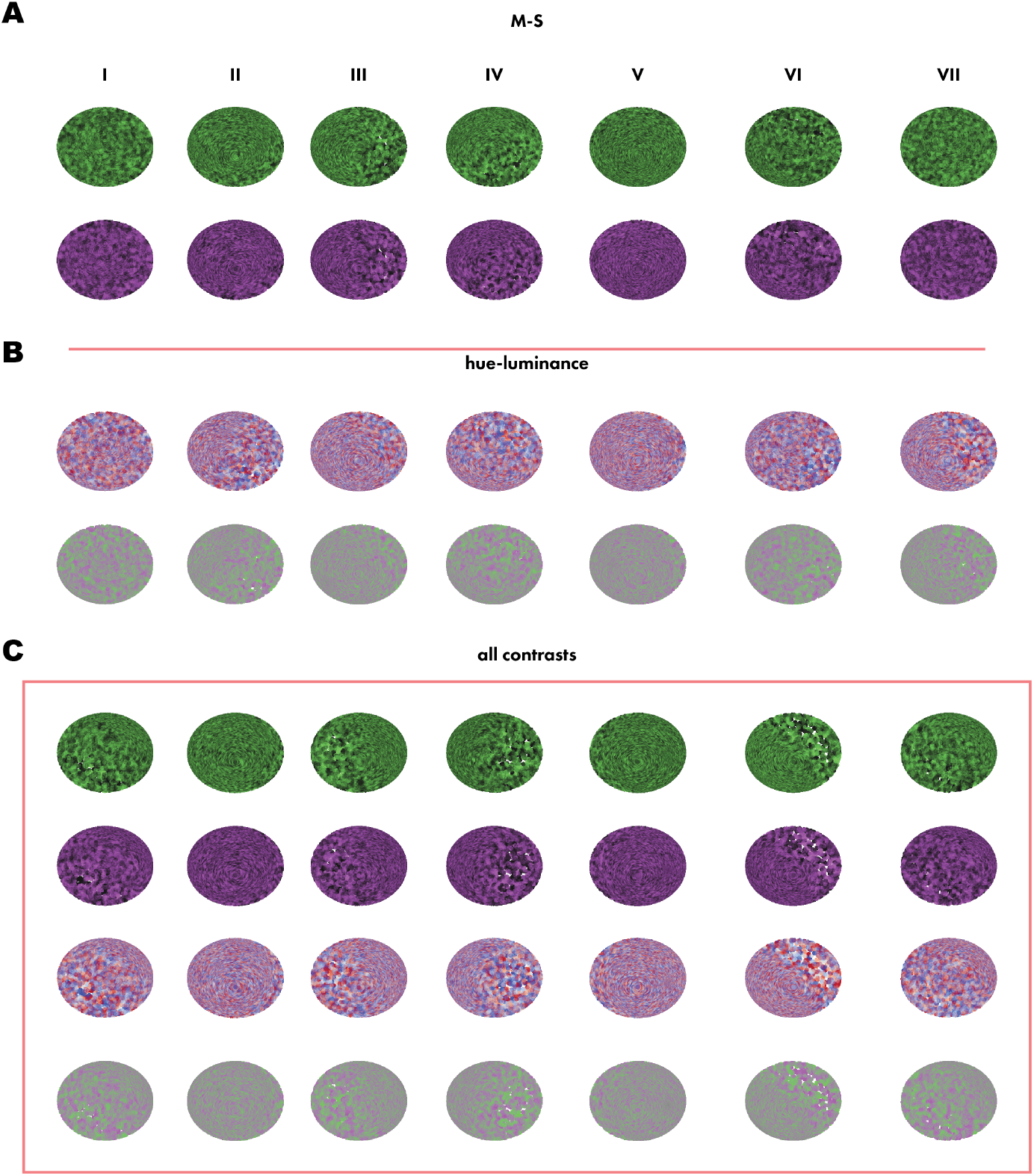
Latent embeddings of V1 population activity for all recordings. Plots of V1 population activity from all recordings (n = 7, I - VII) embedded against (**A**) M-S contrasts, (**B**) hue-luminance contrasts, **C** all contrasts. Recording III was used as the example for Figure 7 and Figure 7 with Figure S8.

**Figure S11:**
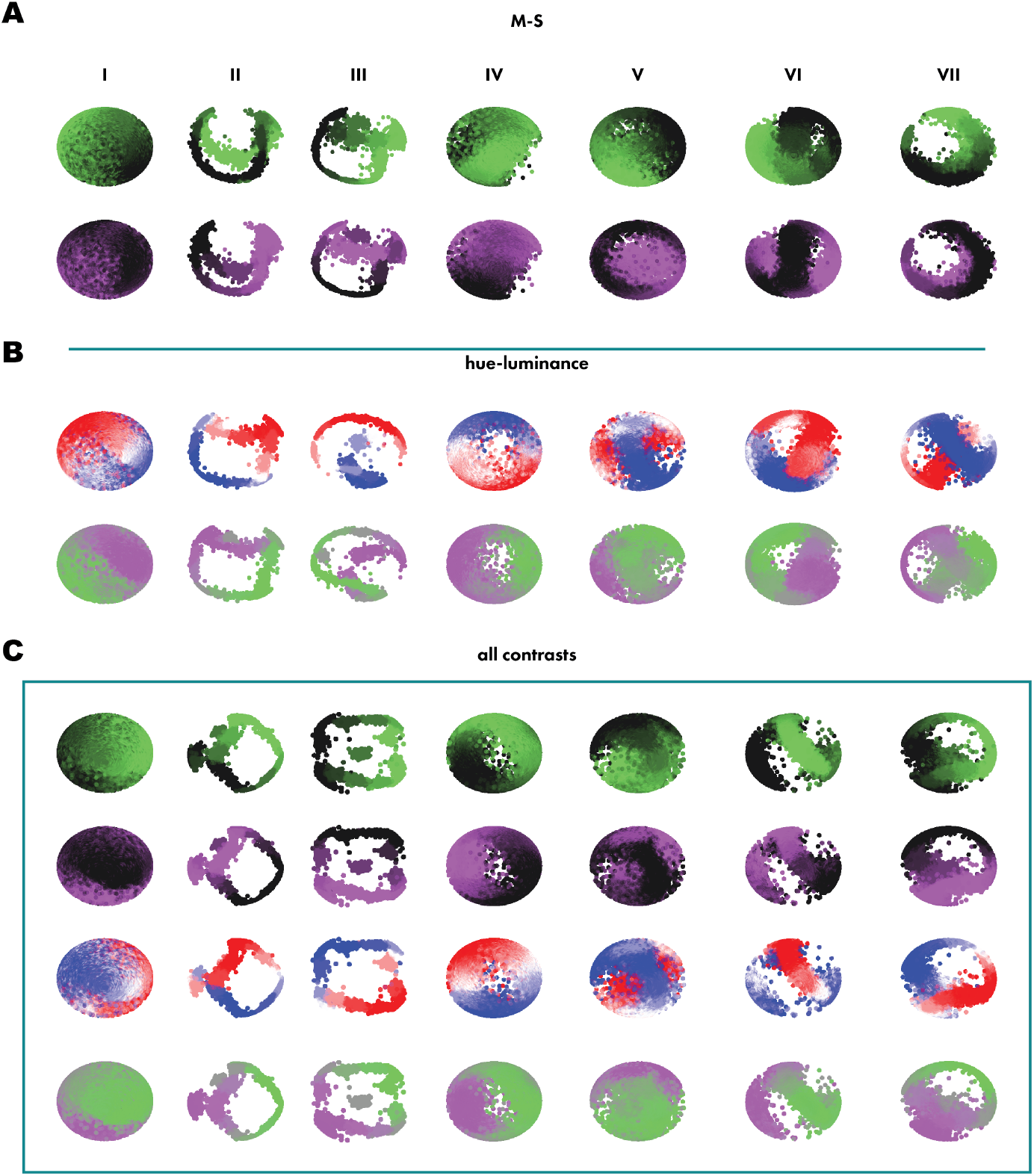
Latent embeddings of trial-shuffled dLGN population activity for all recordings. Plots of trial-shuffled dLGN population activity from all recordings (n = 7, I - VII) embedded against (**A**) M-S contrasts, (**B**) hue-luminance contrasts, (**C**) all contrasts. Recording III was used as the example for Figure 7 and Figure 7 with Figure S8.

**Figure S12:**
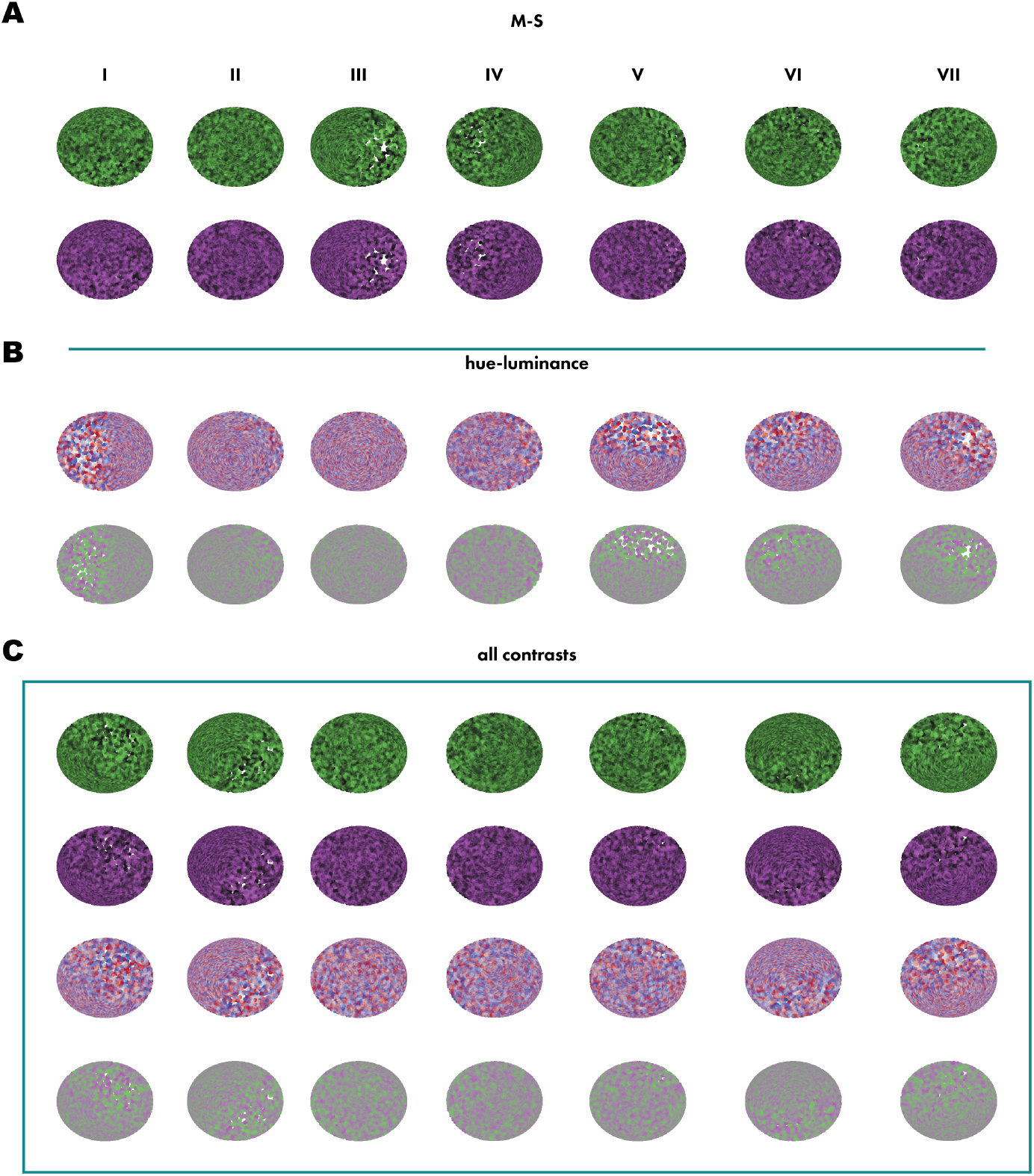
Latent embeddings of shuffled V1 population activity for all recordings. Plots of trial-shuffled V1 population activity from all recordings (n = 7, I - VII) embedded against (**A**) M-S contrasts, (**B**) hue-luminance contrasts, (**C**) all contrasts. Recording III was used as the example for Figure 7 and Figure 7 with Figure S8.

